# An iterative search algorithm to identify oscillatory dynamics in neurophysiological time series

**DOI:** 10.1101/2022.10.30.514422

**Authors:** Amanda M. Beck, Mingjian He, Rodrigo G. Gutierrez, Gladia C. Hotan, Patrick L. Purdon

## Abstract

Neural oscillations have long been recognized for their mechanistic importance in coordinating activity within and between brain circuits. Co-occurring broad-band, non-periodic signals are also ubiquitous in neural data and are thought to reflect the characteristics of population-level neuronal spiking activity. Identifying oscillatory activity distinct from broadband signals is therefore an important, yet surprisingly difficult, problem in neuroscience. Commonly-used bandpass filters produce spurious oscillations when applied to broad-band noise and may be ill-informed by canonical frequency bands. Curve-fitting procedures have been developed to identify peaks in the power spectrum distinct from broadband noise. Unfortunately, these ad hoc methods are prone to overfitting and are difficult to interpret in the absence of generative models to formally represent oscillatory behavior. Here we present a novel method to identify and characterize neural oscillations distinct from broad-band noise. First, we propose a new conceptual construct that makes clear, from a dynamical systems perspective, when oscillations are present or not. We then use this construct to develop generative models for neural oscillations. We show through extensive analyses of simulated and human EEG data that our approach identifies oscillations and their characteristics far more accurately than widely used methods, achieving near-perfect recovery of the number of oscillations when the SNR exceeds a very modest threshold. We also show that our method can automatically identify subject-level variations in frequency to overcome the limitations of fixed canonical frequency bands. Finally we demonstrate how our method can extract clinically-relevant neurophysiological features with greater statistical efficiency other established methods.

## 1 Introduction

Neurophysiological time series such as local field potentials (LFP), electrocorticograms (ECoG), and electroencephalograms (EEG), contain a wealth of information about the properties of the underlying neural systems that generate these data. Oscillations have long been recognized for their mechanistic importance in coordinating activity within and between neural circuits [1]. More recently, oscillatory phase relationships, wave phenomena, and nonsinusoidal quasi-periodic waveforms have been proposed as mechanisms for neural computation [2, 3]. On the other hand, broad-band, non-periodic signals, which are thought to reflect population-level neuronal spiking activity, are also ubiquitous in neural data. The mechanistic significance of this broad band activity is not fully understood and is an area of active investigation. Recently, the spectral slope for such broad-band non-periodic signals has been proposed as a potential marker of excitatory-inhibitory balance [2].

While the distinction between oscillatory and non-periodic dynamics may seem a trivial one that would be obvious by inspection, in practice it can be very difficult to determine whether an oscillation is present distinct from broad-band noise spanning a frequency range of interest. A common approach for analyzing oscillations is to bandpass filter the observed signal in the frequency range of interest and examine the resulting waveforms. Unfortunately, applying a narrow-band filter to a broadband signal will generate a spurious oscillation that merely reflects the oscillatory properties of the filter (Supplementary Figure 1) [4]. Identifying the correct frequency limits for such filtering is a further problem. In neuroscience and clinical neurophysiology we describe oscillations in terms of canonical frequency bands, but neural oscillations in practice do not necessarily fall exactly within those bounds. For instance, as people age, it is well-known that posterior alpha waves can decrease in frequency, sometimes below the canonical 8 to 12 Hz alpha band [5–7] and due to individual variation, oscillations such as sleep spindles, which are generally considered to be between 11-16 Hz or conservatively 12-15 Hz, may fall outside the expected frequency range [8, 9]. To overcome these problems, recent analysis methods have employed curve-fitting procedures to identify peaks in the power spectrum distinct from broadband noise [10–13]. The advantage of these approaches is that they are intuitive and are easy to apply using freely-available software. On the other hand, these methods are ad hoc, lack formal procedures for statistical inference, and rely on parameter tuning that may not be robust and could require significant user-intervention. The main shortcoming of these methods, however, is that the relationship between the observed data and the properties of an underlying dynamical system consistent with these data cannot be ascertained; i.e., there is no generative model that can summarize or explain the key properties of interest. The absence of a generative model impairs not only interpretation, but also statistical efficiency.

Here we present a technical note that describes a novel approach to neural time series analysis that can robustly identify and characterize neural oscillations distinct from broad-band background noise. To achieve this, our method identifies a generative model whose parameters describe the frequency, damping, and gain of an underlying oscillatory system [14] and uses an expectation maximization (EM) algorithm to fit the model parameters [15]. Our method automatically identifies the number of oscillations present in a time series with minimal user intervention and can do so on a single time series that may contain only a few cycles of an oscillation of interest amidst other possibly larger oscillations. Our method achieves this using an iterative procedure that identifies oscillations in order of decreasing power and strips them away, making it possible to focus on a single oscillator at each scale. This iterative procedure greatly improves learning accuracy for both the number of oscillations and their characteristics. Finally, based on our generative model, our method can extract multiple simultaneous oscillatory time series, along with their phases and amplitudes. We demonstrate the performance of our method using both simulated and human resting state EEG data and show that our approach is far more accurate than existing, widely used methods both in terms of the number of oscillations identified and the estimated oscillation waveforms. We also illustrate how our method can identify individual subject-level parameters for oscillations that can overcome the limitations of fixed canonical frequency limits.

## 2 Methods

### 2.1 Representing oscillatory dynamics using linear dynamical systems

In previous work, oscillatory structure has been identified by characterizing the shape of the spectrum, with the rationale that an oscillatory signal should necessarily show a peak in the spectrum corresponding to each oscillation [10– 12]. We take a slightly different approach: we propose that an oscillation is present if the underlying dynamical system that generated the observed signal is oscillatory by nature. A linear dynamical system, such as:

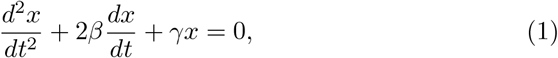

may oscillate if and only if the roots of the characteristic equation are complex. In this case, the (deterministic) waveforms that may be generated by this system take the form

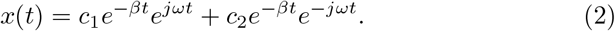

If we regard the observed neurophysiological signal as the real part of *x*(*t*) we can set *c*_2_ = 0.

We can evaluate the continuous-time solution above at discrete intervals of Δ*t* to represent sampled data:

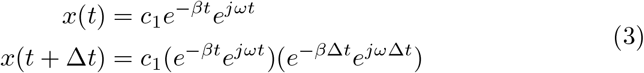

If we separate the real and imaginary components for Equation 3 we can see that future values of the oscillatory state are related to past values by a damped rotation matrix:

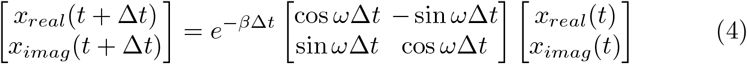

The constant *e*^−*β*Δ*t*^ must be equal to or less than one for the system to be stable. In this scenario, in the absence of any additional external inputs, *x*(*t*) will decay to zero in an oscillatory manner like an underdamped spring-mass system [4, 16]. When driven by a external random input, such as an independent, identically-distributed Gaussian noise source, the system will continue to oscillate in a stochastic manner. The oscillatory properties of nonlinear systems can be analyzed in a similar way by employing a linear approximation in the neighborhood of the system’s fixed points [17, 18]. We will therefore focus on linear systems, with the understanding that the method can be adapted to nonlinear systems.

Given this discrete-time approximation for the continuous-time solution to a linear oscillatory system, we can construct a discrete-time state space model to describe how the system evolves in time when driven by stochastic inputs:

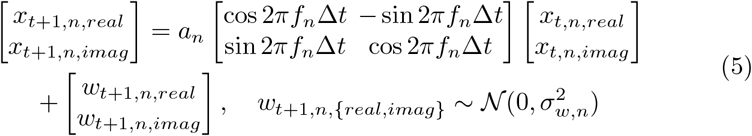

where *n* is the label for a particular latent oscillator, ***w***_*t,n*_ is the Gaussian state noise, *ω* = 2*πf* for *f* in Hz and *a* = *e*^−*β*Δ*t*^. We note that in the frequency domain, *a* is the radius of the complex pole for this system. We can represent a single oscillator *n* more concisely in matrix vector notation:

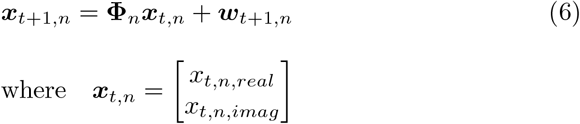

where 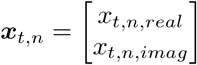

This model of oscillatory dynamics was first proposed by Wiener (1966) and later re-discovered by Matsuda and Komaki (2017) and makes it possible to represent oscillations that are influenced by random perturbations in frequency and amplitude [14, 19].

We can expand this state space model to accommodate multiple simultaneous oscillators at different frequencies:

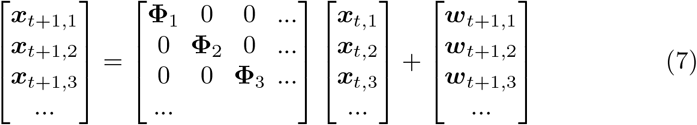

which we represent more concisely by:

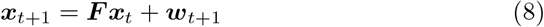

where ***x***_*t,n*_ ∈ ℝ ^2^ and ***x***_*t*_ ∈ ℝ^2*N*^ where *N* is the total number of oscillations. Note that we consider each oscillator to be independent of the others. Finally, we represent the measured neurophysiological signal as a linear combination of the real components of the multiple latent oscillators and Gaussian observation noise, *r*_*t*_:

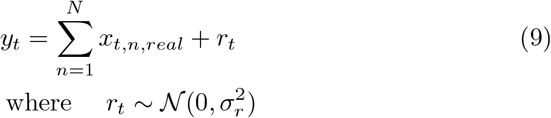

or, equivalently and more concisely:

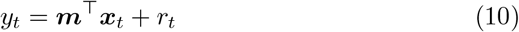

where ***m*** = [1 0 1 0 …] ^*T*^ and *y*_*t*_ is the recorded data at a single electrode at time *t*.

### 2.2 Interpreting the State Space Model as a Filter Informed by the Data

Contrary to bandpass filtering with fixed or arbitrary frequency bands, with this model it is possible to estimate the relevant oscillatory components directly from the time domain data. We note that the frequency response for each oscillator component has “soft” limits consistent with a second-order physical system as described in Equation 1. In contrast, bandpass filters are typically constructed with sharp band limits that are achieved using high-order linear discrete-time systems that can create spurious oscillations when applied to broad band signals (Supplementary Figure 1) [10, 20]. The form of this model effectively leads us to learn a data-driven filter for each oscillator component, with varying bandwidth and shape, using the expectation maximization algorithm. The fitted parameter *f*_*n*_ shows the center frequency of oscillation *n*, while *a*_*n*_ controls the shape and bandwidth, and 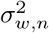 denotes the power or variance assigned to this oscillation.

The challenge with this model is that the parameter estimation problem can be highly non-convex, particularly when an oscillation is small compared to neighboring oscillations or observation noise. Model selection for the correct number of oscillators is also challenging, and again further complicated when oscillations are small compared to other signals or noise. To overcome these challenges, we developed an iterative algorithm that identifies the number of oscillations present in a signal while estimating all model parameters. This method introduces priors to help constrain parameters such as the oscillator frequency and noise covariances that are not robustly estimated otherwise. It also employs a novel initialization procedure that uses prediction errors from lower-order models to identify and initialize missing components in a putative higher-order model.

### 2.3 Iterative Oscillator Component Algorithm (iOsc)

In our experience, although the method described in Matsuda and Komaki (2017) could identify relevant oscillatory signal components, the method could not reliably recover the correct number of oscillators, tending instead to identify too many components. This seemed to occur because the method significantly under-estimated the observation noise variance, which then led to the identification of spurious oscillators at high frequencies to compensate. When we tried to restrict the number of components, we found that the larger oscillatory components could be overfitted with multiple oscillators at the expense of smaller neighbors, particularly in scenarios where the oscillations spanned a wide dynamic range, as is often the case in neuroscience data. To overcome these challenges, we designed an algorithm to first fit the highest power oscillations and then use the unexplained power in the observed signal to initialize additional putative oscillators and relevant hyperparameters. We used a von Mises prior on the oscillation frequency to constrain the estimated frequency of each additional oscillation (Appendix C.1). We also used an inverse gamma prior on the observation noise variance to prevent it from decreasing too much (Appendix C.2). Appendix D describes the complete data likelihood for the model after the addition of these priors.

Here we introduce notation to distinguish the parameters for different models *ℳ*_*i*_ : the parameter 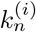 refers to the parameter *k* for oscillation *n* in a model containing *i* oscillations (also referred to as iteration *i*). The procedure for initializing the very first oscillator is described in Appendix B. It is structurally identical to initialization of subsequent iterations except that the observed data are used in place of the prediction error. The following describes the iterative process of fitting subsequent oscillations in the model.

#### 2.3.1 Measuring model error with the one-step prediction error

In order to fit the *i*-th oscillator, we wish to examine any remaining oscillatory information not represented by the (*i* − 1) oscillator components in the previous model.

The residual error–i.e., the difference between the observed data and the estimated signal after filtering or smoothing–would seem to be a natural choice for identifying any remaining oscillatory structure poorly captured by the model. However, at each time step *t* the filter and smoother update the estimate of the hidden states 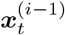 using the observed data at time *t*, essentially compensating for deficiencies in the model-based prediction. Accordingly, in practice we found that the residual error for the filter and smoother contained little information on remaining oscillatory structure.

Alternatively, the one-step prediction error (OSPE) 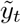, also referred to as the innovations [15, 21], relies on the predictions of the latent states at the next time point using only the model and no additional observed data:

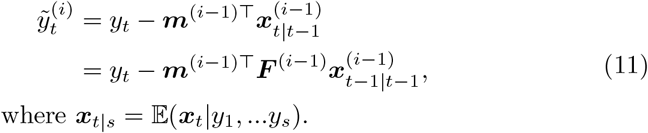

Where ***x***_*t*|*s*_ = 𝔼 (***x***_*t*_|*y*_1_, …*y*_s_).

We found that any remaining structure not accounted for in the (*i* − 1)-th model was visible in the OSPE. We therefore used the the OSPE to initialize the parameters for the *i*-th oscillator.

We use a low-order autoregressive (AR) spectral estimate as described in equation 12 to analyze the structure of the OSPE,

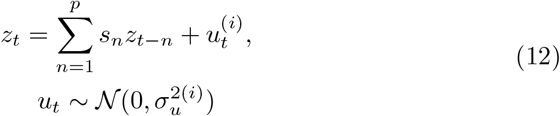

setting 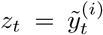 and *p* ∈ *{*7, 9, 11, 13*}* in order to represent a small number of oscillations and one random walk component. We use the Yule-Walker algorithm to obtain the parameters for this AR model.

#### 2.3.2 Parameter Fitting

We initialize the oscillator frequency of the *i*-th oscillator component, 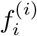, and the mode of the von Mises prior on this component’s frequency using the frequency of the largest magnitude pole in the AR model (Appendix B.1). For this *i*-th oscillator component, we initialize the radius 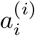 at the radius of the pole. We initialize the state noise variance, 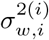, the observation noise variance, 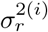, and the mode of the inverse gamma prior for the observation nose variance to 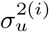. We then use an EM algorithm to fit the parameters for all oscillators in the model. We provide the update equations in Appendix E.

Figure 1 illustrates this iterative process over multiple components. Figure 1a shows the resulting model after fitting a single oscillator component to data that clearly has two peaks. Figure 1b shows a peak in the power spectrum of the OSPE, showing that the alpha power is not well modeled by the oscillator fit in the panel above it. The AR(13) fit to the OSPE is shown as a blue dased line and the largest magnitude pole location is shown in green. We can see by eye in Figure 1c that two oscillators seem to match the data well, and the flat OSPE and its corresponding AR(13) fit shown in Figure 1d confirm that the two oscillator model accurately represents the oscillatory structure in the data. The magnitude of the largest pole at this step is also much smaller than that of the pole in the previous step. Supplementary Figure 2 shows that when the von Mises prior is omitted, the second oscillator is pulled towards the slow peak. After two iterations without the von Mises prior, the OSPE shows peaks in power in both slow and alpha frequencies, implying that the fitted model is not capturing all the structure in the data despite having the correct model order.

**Fig. 1:**
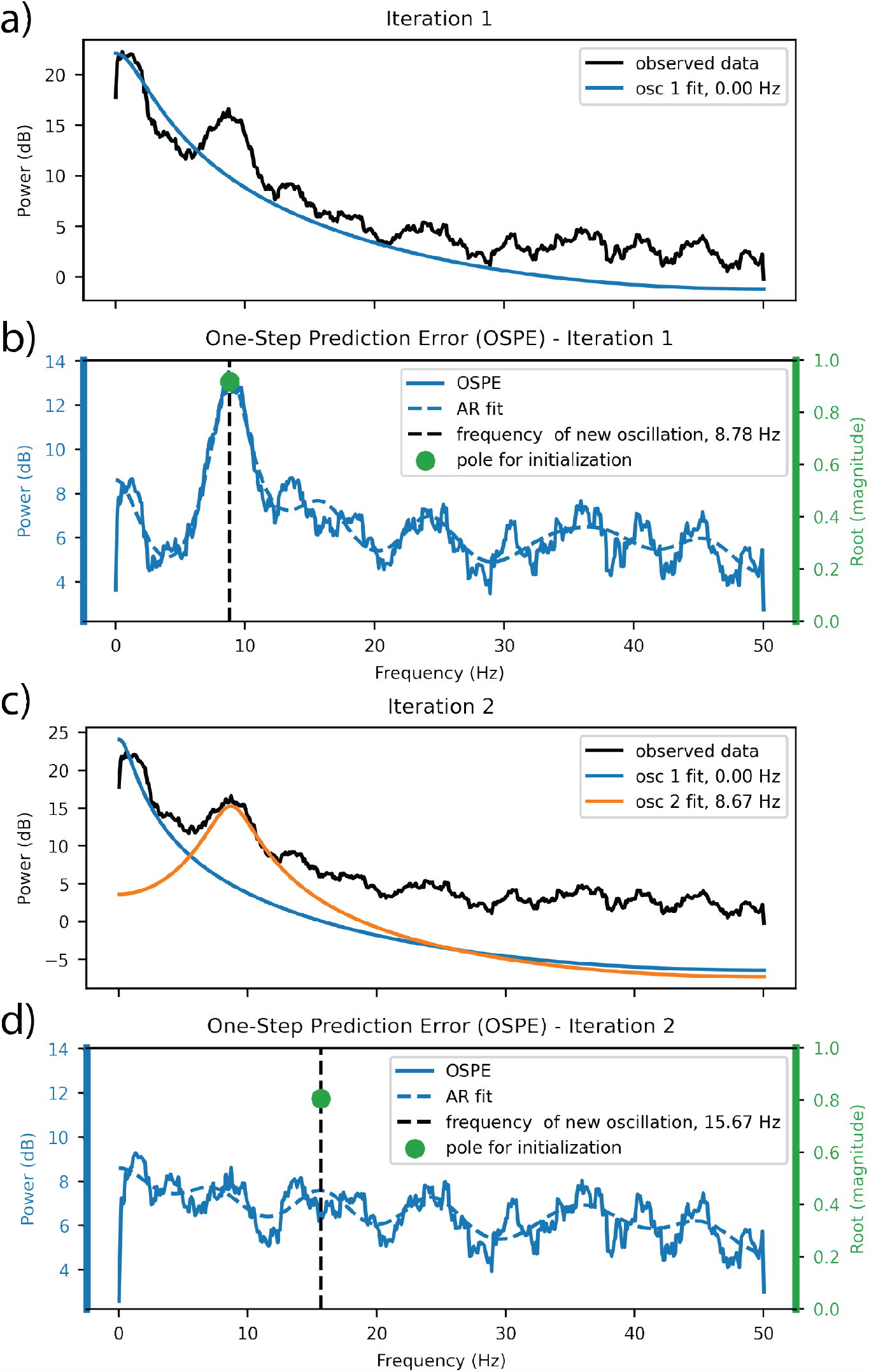
Example of how OSPE shows which portions of the signal are or are not represented by the intermediate model. Simulated data with slow frequency 1 Hz and alpha frequency 8 Hz.

As our model converges to the true model of the system, the OSPE will converge to the observation noise, which we assume to be white (See Appendix F). If the OSPE is white noise, then the best fitting AR(*p*) system will have no autoregressive structure and the autoregressive coefficients will converge to zero, with the variance of the driving noise approaching the variance of the observation noise. Based on this convergence behavior, we will use the AR(*p*) noise variance 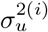 as an approximation of observation noise variance 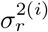, setting the mode of the inverse gamma prior to be 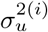.

### 2.4 Model Selection

Model selection criteria such as the Akaike Information Criterion (AIC) and the Bayesian Information Criterion (BIC) have been widely applied to time series modeling problems, including the present class of state space oscillator models as originally described by Matsuda and Komaki. However, we have found that when dealing with state space models at high sampling rates, as is often the case in neurophysiological data, AIC tends to bias model selection in favor of more complex models. This occurs because the log likelihood grows with increasing data length whereas the model complexity penalty, equal to 2*p* where *p* is the number of parameters, remains constant irrespective of the data length (see Appendix H). BIC’s model complexity penalty, *p* ln(*T* ), increases with data length, but in practice tends to favor more complex models as well. Given the prior knowledge that neural recordings tend to contain a small number of oscillations that have different mechanistic origins and functional properties, we prefer a lower dimensional model that may be more interpretable. Although each additional oscillator component should decrease the variance of the unexplained signal, and thus increase the log likelihood, the marginal increase in log likelihood tends to diminish beyond some point. To take advantage of this decreasing marginal change in the log likelihood, as an alternative to AIC or BIC, we used the “kneedle” method [22, 23], which uses the curvature of the log likelihood sequence across the set of models to identify the model order beyond which there are diminishing marginal improvements in log likelihood.

### 2.5 Human Data Statement

The resting state EEG data analyzed in Section 4 of this paper was collected from healthy adult volunteers in an study approved by the Mass General Brigham Institutional Review Board (formerly Mass General IRB) in accordance with the relevant regulations. Informed consent was obtained from all participants. The three subjects used in this paper are female and between the ages of 65 and 67. Their data was collected on a 128 channel EEG system (ANT Neuro North America, Philadelphia, PA).

The EEG data analyzed in Section 5 of this paper was collected from older patients undergoing major abdominal surgery at the University of Chile Clinical Hospital (Santiago, Chile). This study was approved by the local Institutional Review Board in accordance with the relevant regulations. Informed consent was obtained from all participants. Thirty-five subjects were included in the analysis. Cognition was evaluated before the surgery with the Montreal Cognitive Assessment Method. Intraoperative EEG data was collected on a 16-channel system (BioSemi, Amsterdam, The Netherlands). All subjects underwent general anesthesia with sevoflurane during the surgery.

## 3 Results: Simulated Data

To investigate the performance of our algorithm (iOsc) and compare it with other methods [10, 14] we constructed simulated data consisting of slow and alpha oscillations from the generative model class using parameters derived from fitting this model to eyes-closed resting state human EEG data. We produced simulations with varying alpha state noise covariances 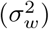 to vary alpha power and SNR, simulating 50 10-second datasets at each state noise covariance value. Figure 3 shows the relative theoretical power spectra of the simulated alpha components compared to the non-alpha components, which consist of the slow oscillation and the observation noise. The theoretical alpha band SNR is then given by the ratio of alpha power to non-alpha power between 8 and 12 Hz. We compare the results of the previously described iterative oscillator algorithm to the Matsuda and Komaki method [14], which uses a similar oscillator structure but a different method of parameter fitting and model selection, as well as a frequency-domain peak fitting algorithm (FOOOF) [10]. We characterize the performance of these methods by evaluating the number of oscillations identified as well as the accuracy of the fitted parameters.

**Fig. 2:**
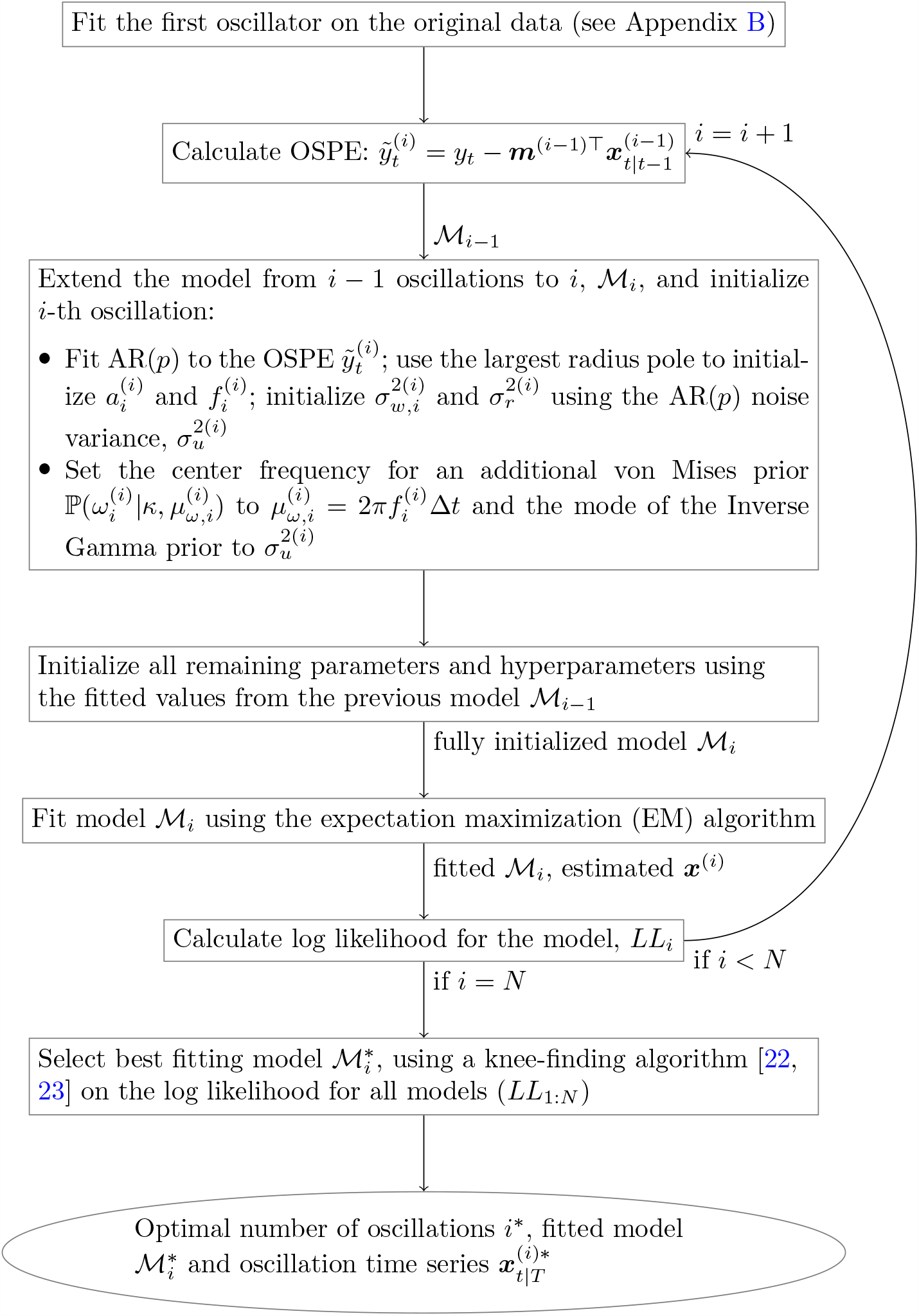
Flowchart describing the proposed iterative algorithm. Note that *N* is the maximum number of oscillations to be considered in the analysis and *T* is the number of samples in the observed data, *y*. The number of AR parameters is a relatively small odd number: *p* ∈ *{*9, 11, 13*}*. Both *N* and *p* can be defined by the user. **Notation:** parameter 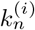 is the parameter *k* for the *n*-th oscillation in a model containing *i* oscillations (also referred to as iteration *i*)

**Fig. 3:**
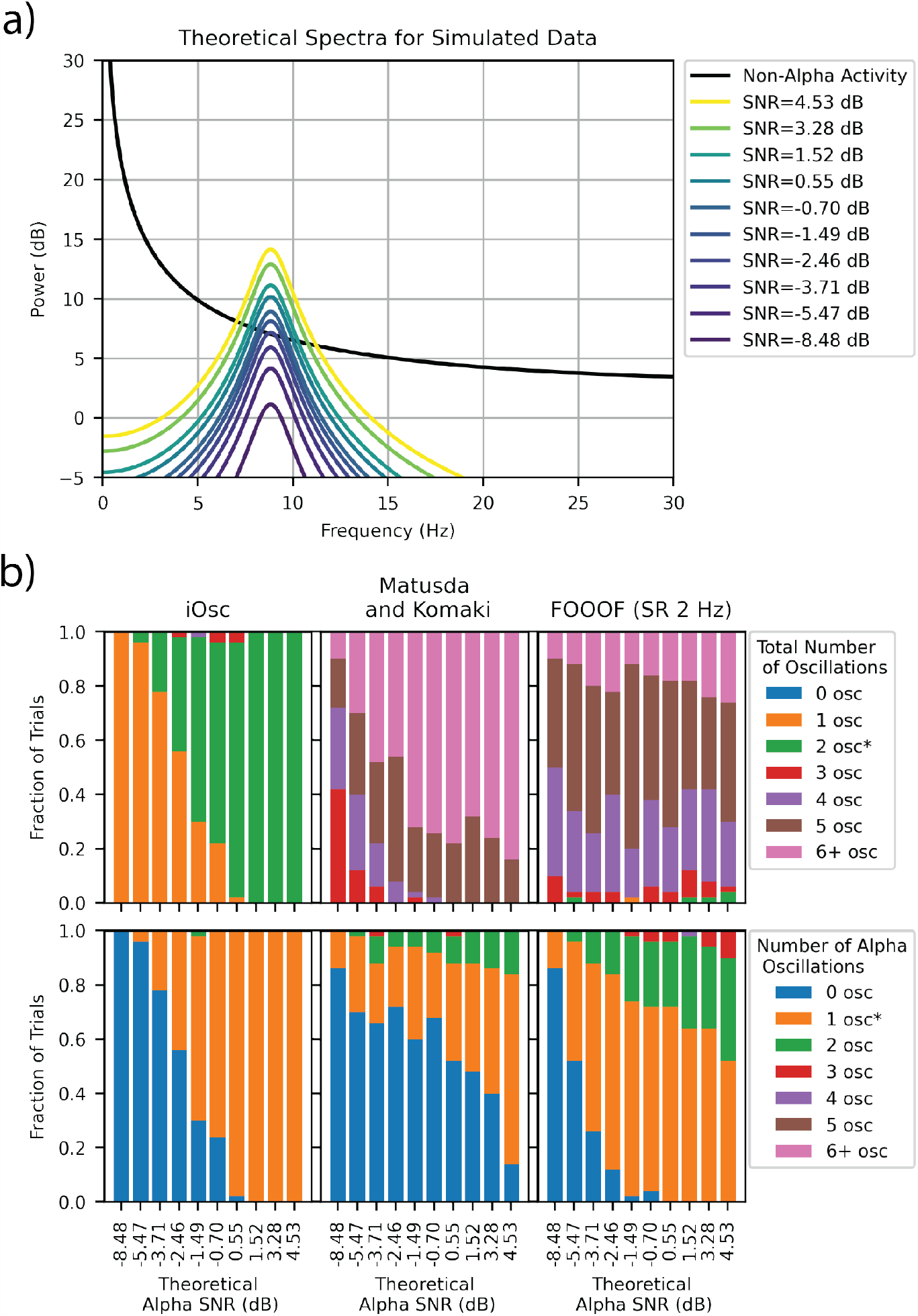
Top: Theoretical Spectra of Alpha and Slow oscillations based on the simulating parameters. Slow parameters: *a* = 0.996, *f* = 0.12 Hz, 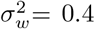, Alpha parameters: *a* = 0.938, *f* = 8.82 Hz, 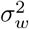 varies to achieve different alphaband SNR values as indicated in the legend. Bottom: Comparing the selected number of oscillations with each method for 50 simulations at each SNR level. The correct number of oscillations is marked in the legend with an asterisk. FOOOF is used on spectra with a 2 Hz spectral resolution.

### 3.1 Identifying the number and frequency of oscillatory components: Comparison with Matsuda and Komaki and FOOOF

#### 3.1.1 Iterative Oscillator Model (iOsc)

Using these simulated data, we can characterize how accurately each algorithm identifies the presence of an alpha oscillation as its size or SNR varies. When a slow oscillation is generated in the absence of an alpha oscillation, iOsc had a false positive rate of 0% across 500 trials. iOsc starts detecting an alpha oscillator above this false positive rate when the alpha oscillation reaches a theoretical SNR of -5.47 dB, as shown in Figure 3. The true positive rate for detecting two oscillations increases as the alpha state covariance and SNR increase, reaching a value of ≥ 68% at an theoretical alpha SNR of -1.49 dB (Figure 3b, top left). The algorithm achieves nearly 100% accuracy at theoretical alpha SNRs of 0.55 dB and higher, incorrectly identifying an extra oscillation a small percentage of the time ( ≤ 6 %).

Figure 4c shows the distribution of the recovered alpha frequencies from the simulated data. For the iOsc model, the 95% confidence interval spans 0.34 Hz and covers the true underlying frequency. In the Supplementary Materials we illustrate the performance of parameter and signal estimation as a function of the alpha range SNR (Supplementary Figures 4 and 5). Overall, as expected, we find that estimates of the alpha oscillation parameters and signal improve with increasing alpha range SNR (Supplementary Figure 4). In addition, we find that the error in the extracted signals appears to be robust to parameter mis-specification, remaining within 5 dB of the true value even at SNRs as low as -0.70 dB where the alpha oscillator parameters remain imperfectly estimated (Supplementary Figure 6).

**Fig. 4:**
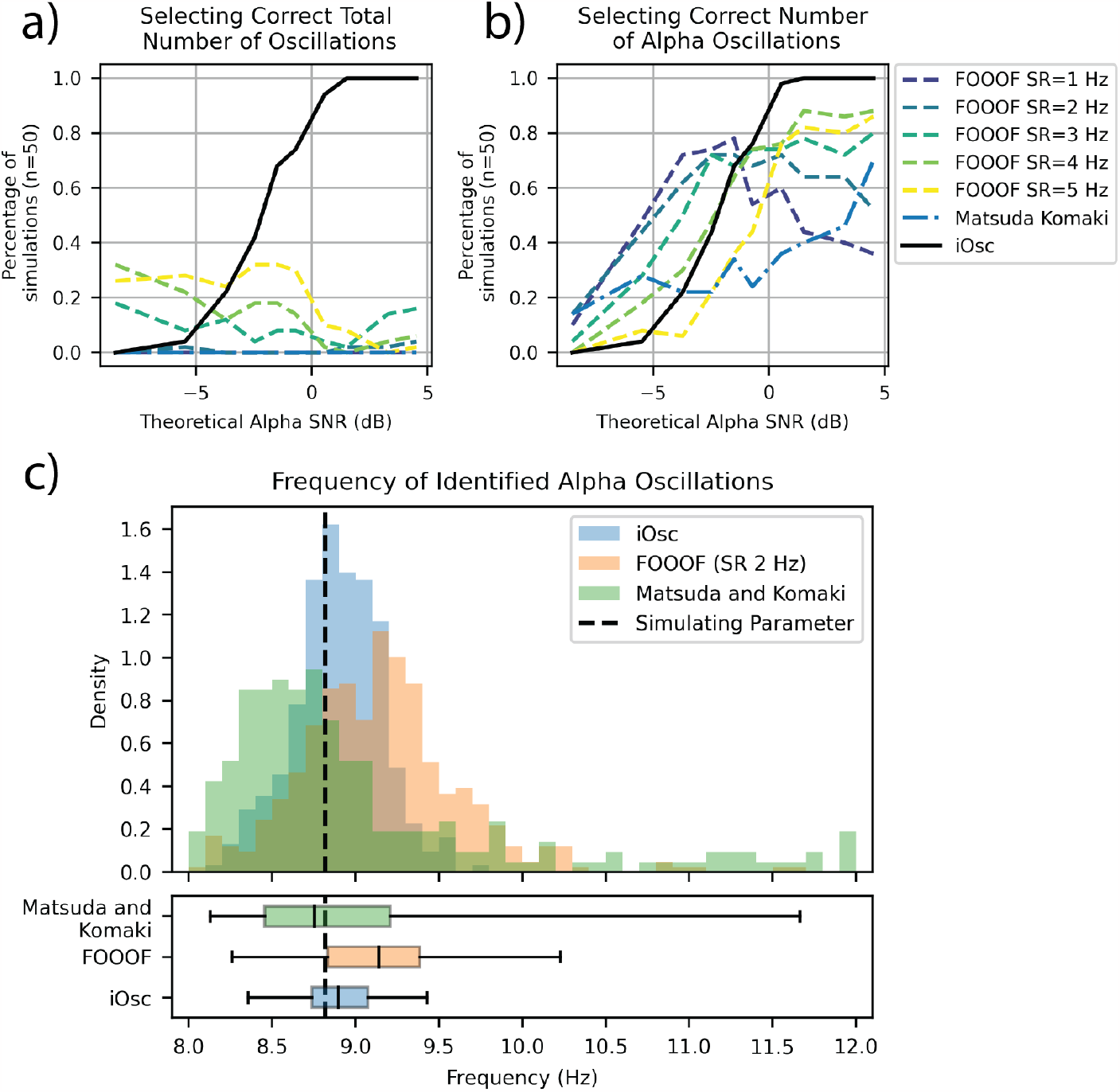
Comparing the number of simulations that have selected the correct number of simulated oscillations. The FOOOF method depends on the spectral resolution (SR) of the signal’s frequency spectrum.

#### 3.1.2 Matsuda and Komaki

In contrast we find that the Matsuda and Komaki algorithm tends to identify too many oscillations (Figure 3b, top middle). In Matsuda and Komaki, the user must first specify *a priori* the maximum number of candidate oscillators, from which AIC is used to select a final model. It is well known that AIC can lead to overfitting of AR models, and we have shown here how this is due in part to the inability of the model complexity penalty to account for increasing data length (see Appendix H). This tendency of AIC could lead to selection of an excess number of oscillator components in the Matsuda and Komaki algorithm. In addition, as discussed in Section C.2, we observed that spurious high frequency oscillator components will be identified if the observation noise covariance is under-estimated; this may also be occurring in the Matsuda and Komaki algorithm.

If we restrict our attention to the alpha band (8 to 12 Hz) and characterize only the number of oscillations identified in this band, the Matsuda and Komaki algorithm can adequately recover the correct number of alpha oscillations even at low SNRs of -5.47 dB. However, this comes at the expense of identifying spurious oscillatory components in total, particularly at high frequencies, which may be misleading in the absence of strong prior knowledge that the content in higher frequencies could be ignored. In comparison to our iterative oscillator model, the Matsuda and Komaki algorithm appears to recover the alpha frequency with less accuracy, showing an 95% confidence interval of 0.76 Hz in the distribution of estimated frequencies (Figure 4c).

#### 3.1.3 FOOOF

Donoghue, et al., have proposed a frequency-domain peak fitting algorithm (FOOOF) to identify oscillatory signal components distinct from broadband noise [10]. FOOOF takes a nonparametric spectral estimate as an input and then identifies an oscillation based on the shape of the spectrum– i.e., the presence or absence of a Gaussian component in a linear regression model of the spectrum. The performance of the method is therefore highly dependent on the resolution and variance of the nonparametric spectrum and on how well the Gaussian-shaped model represents the spectrum of an oscillation. We applied FOOOF to nonparametric spectra estimated using the multitaper method across a range of spectral resolutions [24]. We found that a spectral resolution of 2 Hz produced the most accurate performance over the range of SNRs studied without negatively impacting the peak frequency (Figure 4b). Like the Matsuda and Komaki algorithm, FOOOF tended to identify too many overall oscillations (Figure 3b, top right). This was also true of FOOOF when evaluating only the number of oscillations identified in the alpha band (Figure 4b). As the SNR increases, FOOOF’s overall performance does not improve (Figure 3b, top right) and in alpha band tends to degrade as an increasing proportion of spurious alpha oscillations are identified (Figure 4b). In comparison to iOsc, the FOOOF appears to recover the alpha frequency with less accuracy, showing an 95% confidence interval of 0.55 Hz in the distribution of estimated frequencies (Figure 4c).

### 3.2 Estimation of oscillatory waveform and amplitude: Comparison with bandpass filtering

When a narrowband filter is applied to broad-band noise the filtered signal can appear oscillatory, reflecting the properties of the applied filter (Supplementary Figure 1) [4]. While many neurophysiological signals fit within the bounds of canonically-defined frequency limits, oscillations from a given functional system may not fall exactly within those limits, and there can be significant inter-individual differences in frequency for any given type of oscillation. When these oscillations fall on the edges of the traditional frequency band, a narrowband filter can underestimate the power in an oscillation. For example, posterior alpha oscillations decrease with age, as low as 7-8 Hz [25], and would not be captured by a traditional 8-12 Hz bandpass filter. To account for this inherent variation in oscillatory frequencies, our method does not require defining the limits of a frequency band and can learn the frequencies of oscillations present in the signal. Moreover, because the oscillators in our model are second order systems, they implement gradual transitions between frequency bands that do not introduce artifacts due to the frequency response of the filter. In Figure 5 we compare the performance of our method versus bandpass filtering in a simulated signal consisting of a slow oscillation centered at 1 Hz and an alpha-range oscillation centered at 8 Hz. Figure 5 shows clearly that the bandpass filter misses the peak oscillation power. In frequency domain, the bandpass filtering artefactually shifts the peak alpha frequency to a higher frequency (Figure 5a) and reduces the amplitude of the alpha oscillation, visible in both the waveform (Figure 5b) and the demodulated alpha amplitude (Figure 5c). In contrast, our method accurately recovers the alpha oscillation, preserving its apparent peak frequency (Figure 5a) as well as its power and amplitude (Figure 5b and 5c). Our method also consistently recovers the underlying alpha band time series with lower error than the bandpass filter (Figure 5, table). Although the iterative algorithm uses an assumption of stationarity to fit the model parameters, the Kalman filter and fixed-interval smoother account for stochastic fluctuations in the underlying state as well as the observed measurement, and thus can flexibly track a changing or bursting oscillation amplitude (Figure 5b and 5c). Overall we find that our method performs better than bandpass filtering, particularly in the setting of substantial inter-individual variation in oscillatory frequency that may not conform to pre-defined canonical frequency bands.

**Fig. 5:**
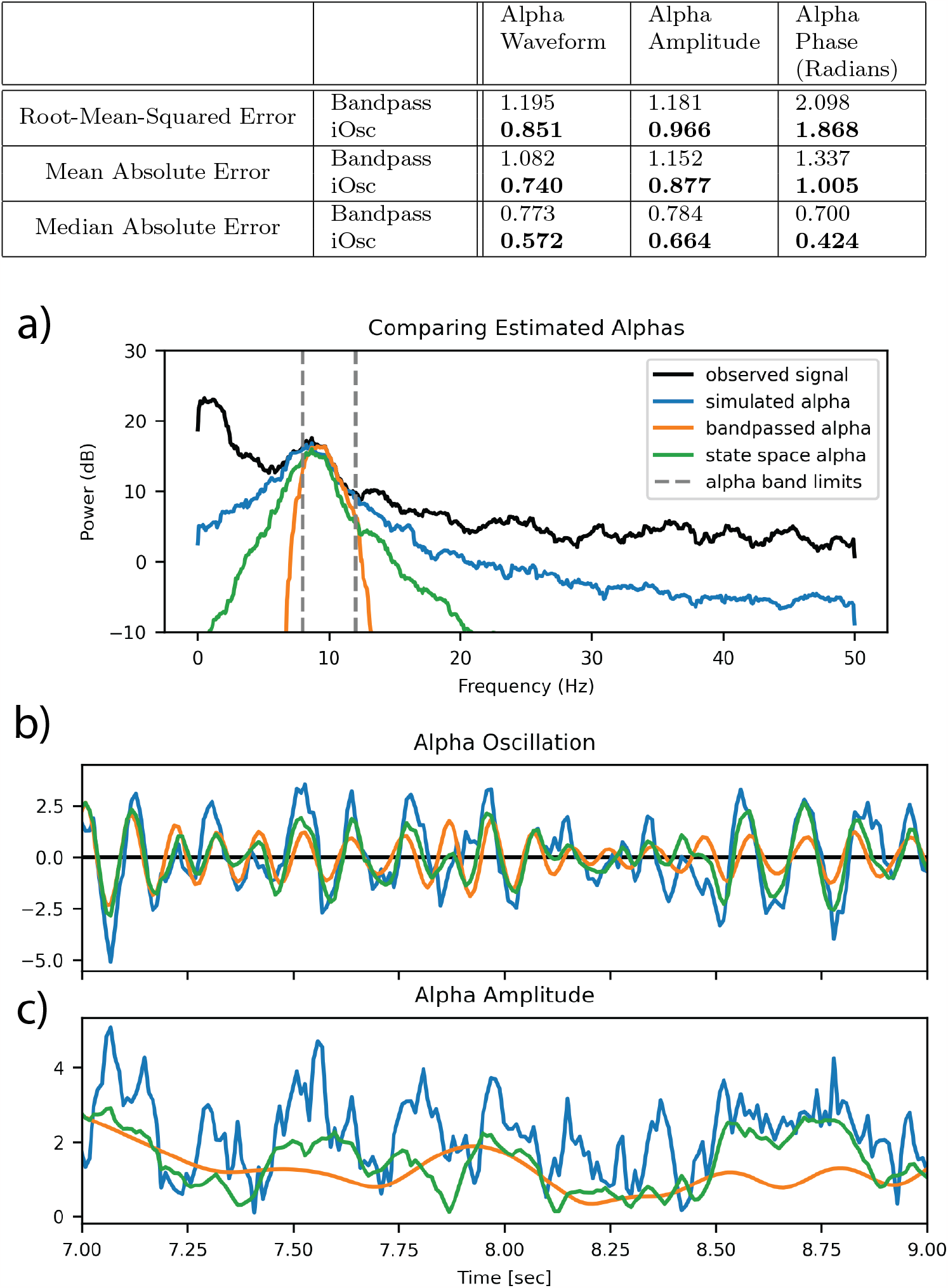
Power captured by the estimated oscillation vs. power captured by bandpass filter (Butterworth bandpass filter IIR order 4 created with scipy.signal, -3 dB points at 8 and 12 Hz and applied forwards and backwards.) Slow param: a=0.97, *σ*^2^=0.4, f= 1 Hz. Alpha param: a=0.9, *σ*^2^=0.5, f= 8 Hz

## 4 Results: Resting State EEG

Here we illustrate how our method applies to real EEG data where the number of oscillations is not known.

We collected data from adult subjects while they were awake with their eyes closed. The occipital alpha oscillations in this state have been extensively studied [26] and may be accompanied by lower-frequency oscillations in the slow- or delta-band [27, 28]. The spectrum in Figure 6a shows an example of 10 seconds of data from a representative subject. We can see that the algorithm identifies a slow and an alpha oscillation with a substantial difference in size of approximately 15 dB. The oscillator model appears to fit the spectrum well (Figure 6a) and also extracts plausible time series within each band (Figure 6c second and third panels). The one-step prediction error time series lacks any obvious oscillatory structure, suggesting that all of the oscillatory activity has been captured in the two model components (Figure 6c, bottom panel).

**Fig. 6:**
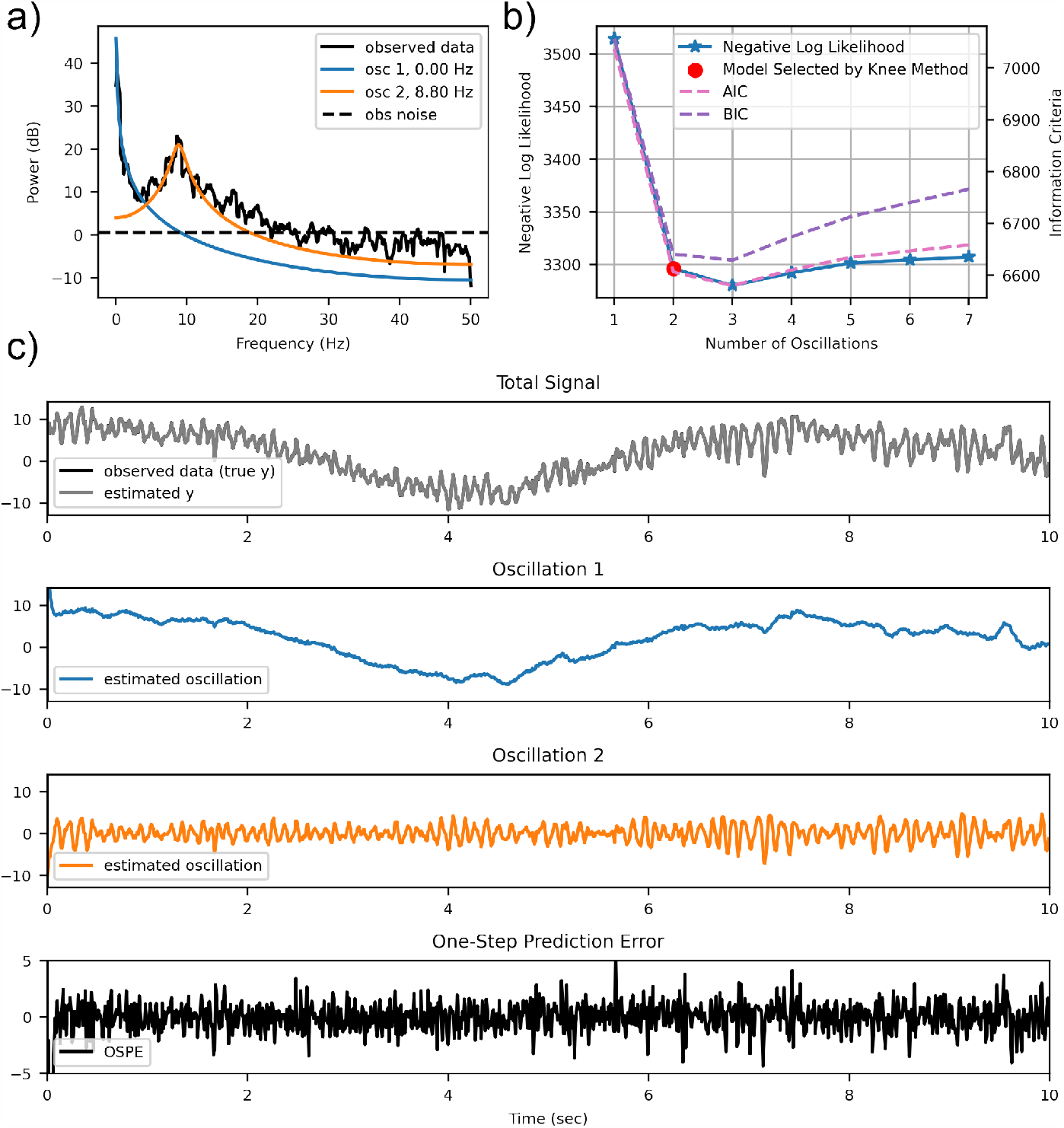
Output of the Iterative Oscillator Algorithm on resting state EEG data. Fitted Slow Parameters: *a* = 0.998, *f* = 0.08, 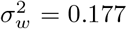. Fitted Alpha Parameters: *a* = 0.946, *f* = 8.82, 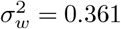. Fitted Observation Noise Parameters: 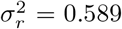

Figure 6b shows the log likelihood, AIC, and BIC for fitted models with different numbers of oscillations for these data. The log likelihood is very similar for models with 2-4 oscillations. The AIC and BIC selection methods would lead us to select three oscillations, but the improvement in the likelhood is very small compared to two oscillations. Selecting the largest likelihood would give us the same result, since the parameter bias is negligible compared to the log likelihood. The knee method selects a model with two oscillations, which corresponds to the point of diminishing marginal returns on the log likelihood. In practice we have found that the knee method tends to identify models with a smaller number of oscillators than AIC or BIC, for reasons discussed earlier and in the Appendix H. In the Supplementary Materials we analyze this data using the Matsuda and Komaki algorithm as well as FOOOF and, consistent with our simulation studies, find that these methods identified a much larger number of oscillations (Supplementary Sections 8 and 9).

In Figure 7 we show how our algorithm can account for significant intersubject variation in the structure of eyes-closed resting state EEG for three female subjects of similar ages. While there are the expected strong slow and alpha range oscillations, we see that shape and power of those oscillations varies significantly between subjects, as does the presence and form of beta oscillations (13-30 Hz). Notice that in Subject 3 our iterative algorithm identifies a model that fits well even though the alpha power is higher than the slow power. This is possible because our algorithm works from the largest oscillations to the smallest, with the von Mises prior acting to prevent the oscillations from moving too much from iteration to iteration. We note that in Subject 3 the beta oscillation (17.47 Hz) is sharp, concentrated around a single frequency, and almost an exact multiple of the alpha oscillation frequency (8.95 Hz). These properties suggest that it could be a harmonic of the alpha oscillation, indicative of a nonlinear oscillation [29, 30] whose structure would not be apparent if analyzed using an alpha-range band pass filter.

**Fig. 7:**
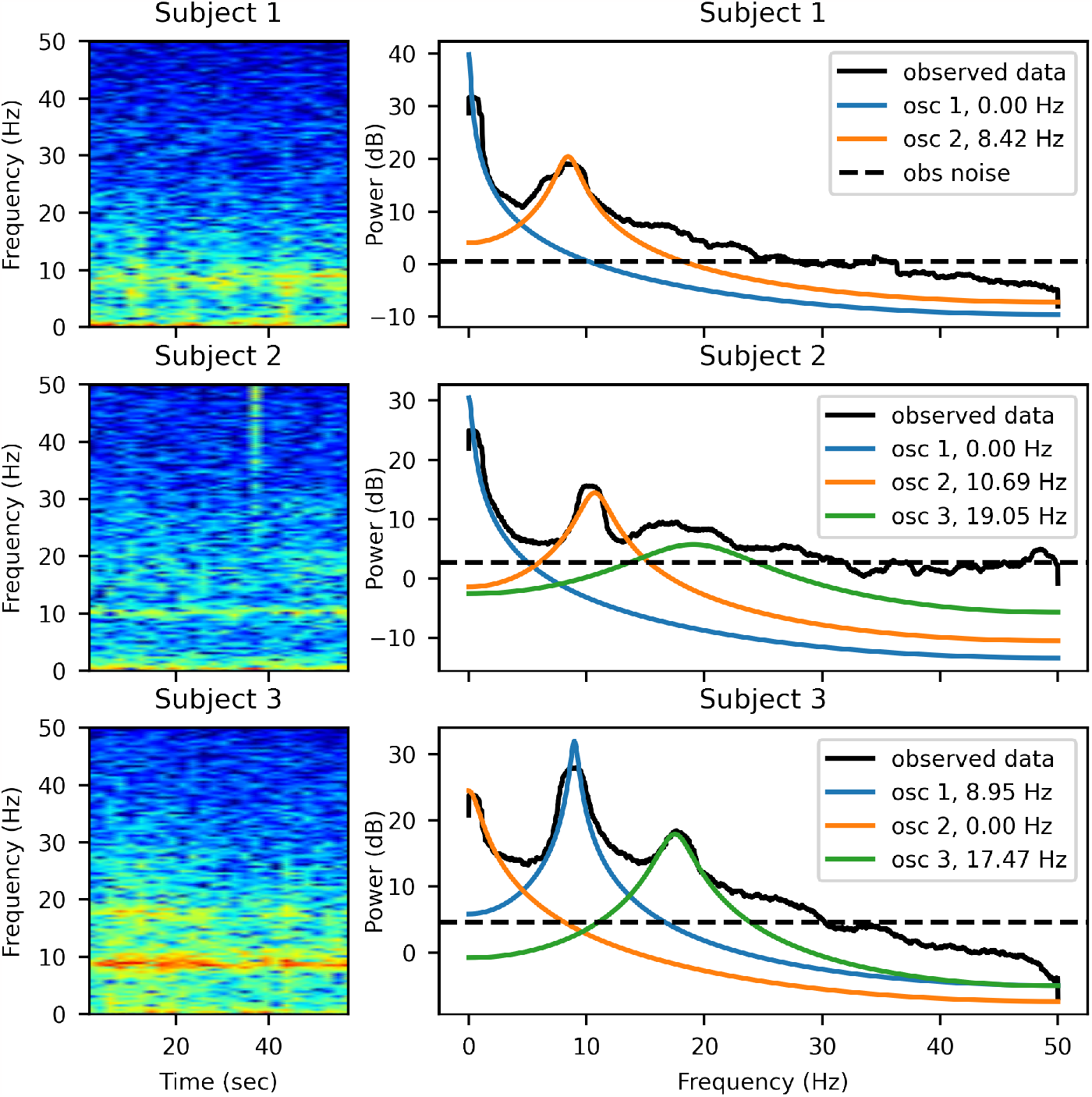
Models selected by iOsc for 60 second EEG segments from three healthy adults of similar ages. All were in resting state with eyes closed.

## 5 Results: EEG Under Anesthesia

Here, in humans under sevoflurane anesthesia, we expect to see slow-delta wave oscillations and alpha range oscillations, with a theta range oscillation at high sevoflurane concentrations [31, 32]. The anesthesia-induced frontal alpha oscillation has already been shown to decrease with declining cognitive health, measured here with MoCA [33, 34], and with age [25]. Our iOsc method consistently identifies 2-4 oscillations (Figure 8, top panel), which is fully consistent with expected behavior of sevoflurane. FOOOF identifies anywhere between 1 to 6 oscillations, which is inconsistent with the known neurophysiology of sevoflurane. In some cases FOOOF identifies only a single oscillation, which likely occurs when FOOOF’s aperiodic model fit captures slow-delta power, exemplifying the difficulty that it has in separating broadband aperiodic activity from oscillatory activity. The Matsuda and Komaki method selects an even wider range of models with up to 14 oscillatory components. This is likely the result of the method’s reliance on AIC to determine the number of oscillations. As illustrated in Supplementary Material Sections 8 and 9, once these oscillation parameters are fit using the EM algorithm, some of them decrease to negligible power, however many spurious oscillations remain.

**Fig. 8:**
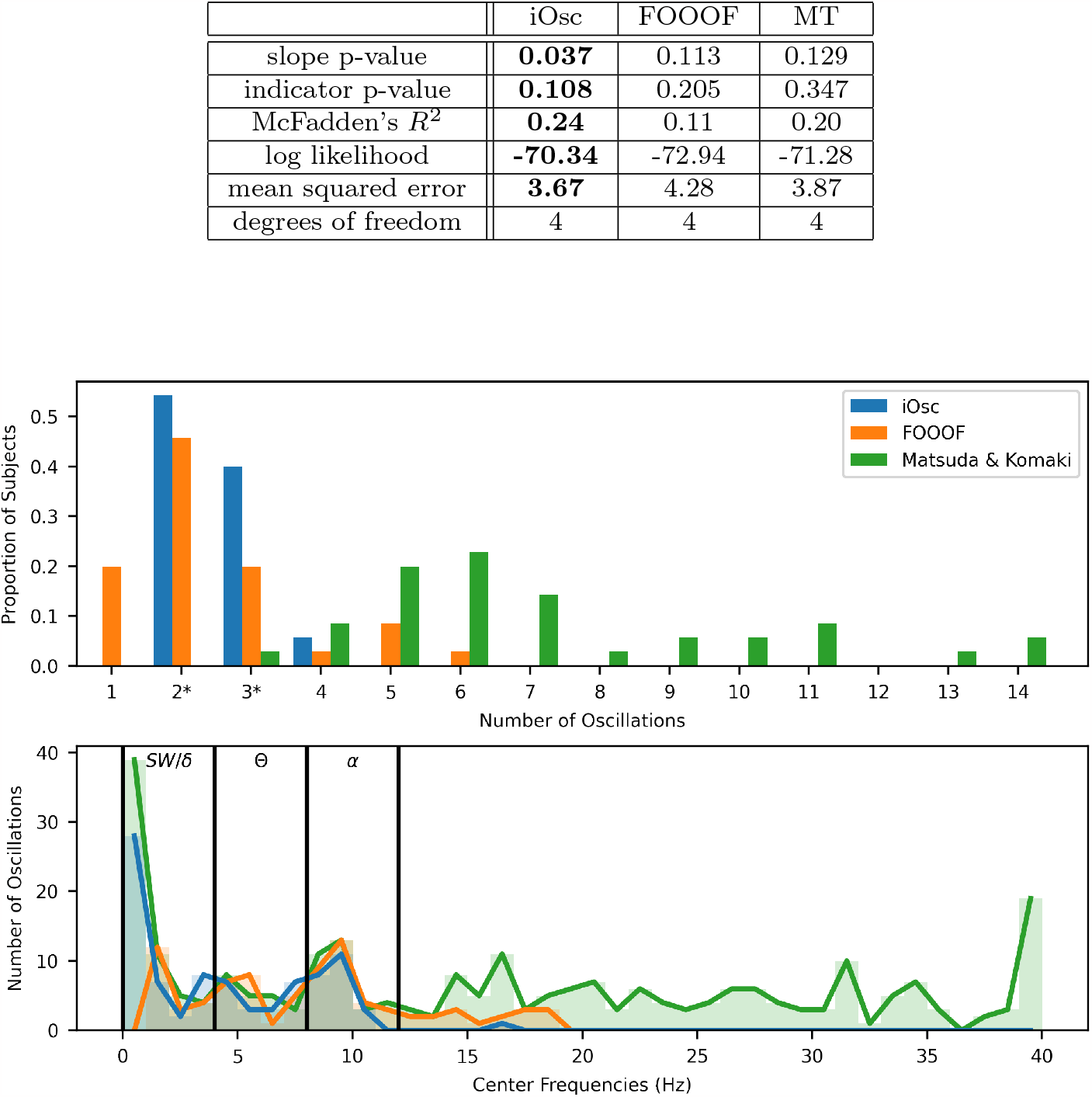
Distribution of models fit to 35 subjects under sevoflurane anesthesia. We expect 2-3 oscillations spread across the slow (*<*1 Hz), theta (4-8 Hz), and alpha (8-12 Hz) frequency ranges.

In Figure 8 (bottom panel), we see the distribution of the center frequencies of the identified oscillations. Center frequencies for iOsc and Matsuda and Komaki are explicitly defined in the model as presented earlier. As expected, we see a large number of slow-delta and alpha range oscillations, with a smaller number of theta range oscillations. FOOOF does not identify any oscillations in the 0-1 Hz range, again due to the confound of its aperiodic component. The range of center frequencies also extends to 19 Hz, which is unexpected for sevoflurane. The Matsuda and Komaki method identifies a high proportion of oscillations in the slow-delta, theta, and alpha bands that are consistent with the iOsc models, but also identifies many spurious oscillations above 10 Hz or at 0 Hz.

We hypothesized that the ability to more accurately identify and parameterize alpha oscillations would make it possible to characterize the relationship between alpha oscillations and cognitive health with greater statistical efficiency. To evaluate this idea, we fit a nested model to describe the presence of each patient’s alpha oscillation and its frequency, if present, as a function of the patient’s MoCA score in n = 35 subjects from the previously reported study described earlier [34]. We used a logistic regression model to represent the presence or absence of an alpha oscillation, in combination with a linear model to relate the alpha frequency to the MoCA score. We compared the performance of features (presence or absence of alpha oscillation, alpha frequency) extracted by iOSC to those from FOOOF as well as multitaper (MT) spectral estimation [24]. For the latter, we used peak finding on a grid search to identify the presence or absence of oscillations. We defined an alpha frequency range of 7-12 Hz which accounts for the fact that older patients may have lower alpha frequencies [25]. For the MT method, if a peak was not found within the boundaries of the alpha band, the alpha oscillation was considered absent. In the case of multiple oscillations within the alpha band, we used the component with the highest peak power.

The table in Figure 8 shows that the slope on the linear portion of the model has a significant p-value only for the iOsc alpha frequency and the iOsc method has the best pseudo-*R*^2^ value. Although the indicator’s p-value is not significant for any method, the iOsc p-value is 47% better than FOOOF and 69% better than MT. The significance of the indicator could be further explored in a larger dataset and may be affected by other factors, like age and amount of sevoflurane. The overall model fit for iOSC is better than FOOOF or MT, as quantified by the log likelihood and mean squared error. The full model fits are shown in Supplementary Figure 16.

## 6 Discussion

Here we present a novel approach for analyzing dynamics in neural signals that can accurately identify the number of underlying oscillations within time series data. We thoroughly characterized the performance of this algorithm using a comprehensive set of simulated and human data where the correct number of oscillations is either known *a priori* or well-characterized. Our simulation studies show that our method can achieve near-perfect recovery of the number of oscillations when the SNR exceeds a very modest threshold. We built upon previous work by Matsuda and Komaki on state space models of neural oscillations [14], but developed a novel iterative search algorithm to improve performance. Compared to existing methods such as FOOOF [10], the state space modeling approach described here makes it possible to extract oscillatory time series as well as the properties of each oscillation, including the frequency, bandwidth, and the instantaneous phase and amplitude, directly from the model. Given the mechanistic importance of oscillatory dynamics in neuroscience, we anticipate that application of this method could significantly improve neural data analyses and reveal novel insights that would not be possible with current approaches.

A major conceptual advance in our approach has been to articulate a more principled answer to the question “What is an oscillation?”. Rather than define an oscillation only in terms of the shape of an observed waveform [3] or spectral estimate [10, 35, 36], we appeal to the fundamental properties of an underlying physical or physiological system that might have generated the observed signal, and recognize that oscillations are possible only in systems where the roots of the characteristic equation are complex. We then show that, under such circumstances, continuous-time oscillations can be represented in sampled data in the state space form proposed initially by Weiner [19] and later re-discovered and further developed by Matsuda and Komaki [14]. Therefore, under our conceptual framework, the model components and parameters employed in data analysis bear a direct relationship to an underlying continuous-time dynamical system. Although our conceptual development and algorithm focuses on linear systems, the same reasoning would apply to nonlinear systems using linear approximations in the neighborhood of the system’s fixed points. Moreover, the generative model class presented here can accommodate nonlinear oscillations by including higher order harmonics [29]. In future work it may be possible to extend the methods developed here to formally characterize nonlinear systems.

A prevailing approach for determining whether oscillations are present in neural time series has been to analyze the shape of the power spectrum [10, 26, 36]. At face value this seems entirely reasonable. However, the dynamical systems perspective we embrace in this work can help us understand why relying only on the power spectrum can lead to disappointing results. In the state space modeling approach used here and in Matsuda and Komaki, the generative model must be able to closely track the observed time domain waveforms, which means that it must reasonably approximate the period (frequency) of putative oscillations as well as their amplitude. In frequency domain, this means that the model must accurately reproduce the power spectrum, but also the *phase response* of the signal. Linear oscillatory systems have a very specific phase response [29], one that is explicitly represented in the oscillator model used in iOSC and Matsuda and Komaki. Methods such as FOOOF, however, focus solely on the shape of the power spectrum and do not account for this phase response. This limits the ability of FOOOF and related methods to distinguish oscillations from similarly-shaped broad-band noise, a behavior that we observed in our analysis of sevoflurane anesthesia (Figure 8). However, for all the advantages of the state space modeling approach, it has a major challenge: a model’s performance depends critically on how well its parameters are estimated and on how well its structure matches the underlying data generating process.

Parameter estimation for state space models can be challenging because the likelihood function is often highly non-convex. We found this to be the case with the state space oscillator model [14]. To make parameter estimation more robust, we introduced prior distributions for the oscillatory frequency parameter (von Mises) and the observation noise (inverse gamma), as well as novel initialization procedures. Identifying the correct number of oscillations, even in simulated data employing the correct data generating process, also proved challenging. Inspired by compressed sensing algorithms such as CoSaMP [37] that identify sparse representations for signals, we developed an iterative search algorithm for state space models that could add oscillatory components based on the residual structure within an error signal. Crucially, we used the onestep prediction error to identify putative new components, recognizing that filtering or smoothing tends to mask deficiencies in the model since they are able to update estimates based on the current or future data, respectively. Our iterative approach is also similar to the Levinson-Durbin recursion [20] for fitting AR models, although we represent the autoregressive dynamics as hidden states in a seasonal adjustment form [21] as proposed by Matsuda and Komaki [14]. Finally, for model selection, we recognized that in AIC and BIC the parameter penalty is negligible compared to the log likelihood (Appendix H), particularly in neurophysiological data with a high sampling rate, leading to a bias towards overly complex models. To account for this, we employed a model selection criteria based on diminishing improvements in log likelihood [22, 23], which allowed our method to consistently select the correct number of oscillations. The fact that our iOSC method can recover the number of oscillations and their underlying frequencies more accurately than Matsuda and Komaki, despite sharing the same underlying generative model, illustrates the value of iOSC’s novel algorithmic contributions.

In this work we have not directly addressed the issue of “1/f” aperiodic dynamics, an area of great interest that may have important mechanistic implications [2]. Instead, we have focused here on maximizing the accuracy with which we can detect, characterize, and extract oscillatory signals. Accurate assessment of oscillatory dynamics has direct bearing on the characterization of aperiodic structure since errors in the oscillatory analysis could clearly propagate and confound the aperiodic analysis. We note that signals derived from oscillatory systems driven by stochastic inputs have an intrinsic broadband character that may overlap with and possibly confound the assessment of non-oscillatory “1/f” dynamics. Similarly, it has been shown that when multiple damped oscillations are combined, their overall spectrum can have a “1/f” shape [38]. These characteristics emphasize how important it is to accurately identify oscillations distinct from aperiodic dynamics. As discussed above, oscillatory systems have very specific phase characteristics, the absence of which has been used by some authors to help identify aperiodic components [36]. Even if oscillations are properly handled, “1/f” properties such as the exponent (or slope on a log-log scale) may be sensitive to a number of confounds or errors including the frequency response of the recording hardware, electromyogram artifacts, and non-neural measurement errors that may have a “1/f” character, to name a few. Although caution is clearly warranted given these challenges, we are optimistic that improved methods could be developed to accurately characterize aperiodic neural dynamics, and remain enthusiastic about the potential mechanistic insights [2] if such efforts were successful.

In conclusion, through this work we hope to advance the important effort to understand oscillatory dynamics in neuroscience. We have introduced a new conceptual construct for characterizing oscillatory neural signals, embodied in a new algorithm that can identify oscillatory dynamics with greater accuracy than current methods. The potential applications in neuroscience are numerous.

## Supplementary information

Please see attached Supplementary Materials.

## Acknowledgments

The authors would like to thank Proloy Das, Carol Wilkinson, Ran Liu, and the rest of the Purdon Lab for their helpful and insightful discussions. Supported by the NSF (GFRP to AMB) and NIH (1R01AG054081 to PLP.).

## Data Availability

Simulated data can be recreated based on the provided code. Human EEG data segments used in this paper can be provided upon request.

## Code Availability

Code for this new method is provided at (https://github.com/mh105/somata) as part of the State-space Oscillator Modeling And Time-series Analysis (SOMATA) package and depends on open source python packages such as kneedle [22, 23] and spectrum [39]. Code used to evaluate the Matsuda and Komaki [14] and FOOOF [10] methods will be provided in Supplementary Materials upon publication.

## Author Contributions

AMB and PLP designed the algorithm. AMB developed and tested the algorithm, and wrote the iOSC portion of the Statespace Oscillator Modeling And Time-series Analysis (SOMATA) package. MH provided critical code review for the MAP estimate components of the iOSC algorithm and designed the SOMATA object structures used to implement iOSC. RG collected and analyzed the sevoflurane anesthesia data. AMB and GH derived the continuous time to discrete time expressions for the generative model.

## Appendix A Terminology and variable definitions

Note:

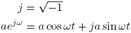

where a = signal magnitude and *ω* = signal frequency in radians.

### Oscillator component

the state space model for of a single oscillation (see Equation 5).

### Oscillation

the time series for a single oscillation (the real part of the latent states for a single oscillator component.)

### Theoretical Spectrum

the analytical spectrum calculated using the model and parameters for a single oscillator component.

### Empirical Spectrum

the multitaper spectrum of the real valued time series from a single oscillation [24].

### Oscillator Model

The state space model including the observation equation. This may include one or more oscillators. An oscillator model is produced after every step of the iterative algorithm.

### Pole Radius

Magnitude of a complex or real root of the AR model. Also known as the **damping factor**. (see Appendix B)

### Spectral Resolution

A property of a spectral estimation method that defines the smallest difference in frequency for which two oscillations may be discerned.

## Appendix B Initialization and parameter estimation for the first oscillation

We begin by fitting a model with a single oscillator. We first fit a low-order autoregressive (AR) spectral estimate to the observed data to initialize parameter estimation for this first oscillator model (*i* = 1):

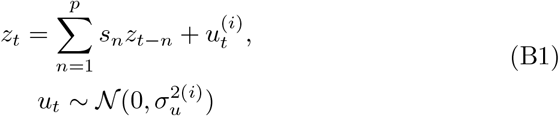

where *z*_*t*_ = *y*_*t*_ and *p* ∈ {11, 13} in order to represent a small number of oscillations and one random walk component. We use the Yule Walker algorithm to obtain the parameters *s*_*n*_ for this AR model. We then use an EM algorithm to fit the parameters for the oscillator model, initializing based on the AR model parameters. We provide the update equations in Appendix E. Here we introduce notation to distinguish the parameters for different models ℳ_*i*_: the parameter 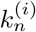 refers to the parameter *k* for the *n*-th oscillation in a model containing *i* oscillations (also referred to as iteration *i*). We initialize the first oscillator frequency, 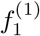, using the frequency of the largest magnitude pole in the in the autoregressive model. For this first oscillator component, we initialize the radius 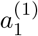 at a fixed, empirically-determined value, *e*^−4.04Δ*t*^, that depends on the sampling interval Δ*t*. We initialize 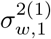 and 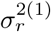 at 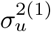. We center the von Mises prior at 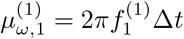.

### B.1 Poles of an AR model

The poles of this AR model define the peaks and valleys of the model’s resulting frequency spectrum [20]. The poles are equivalent to the roots of the characteristic equation, *ξ*_*n*_, and can be computed from the AR coefficients, as seen in Equation B2. If the pole is a complex number, then it has a radius equal to its magnitude and a frequency equal to its phase scaled by the sampling frequency.

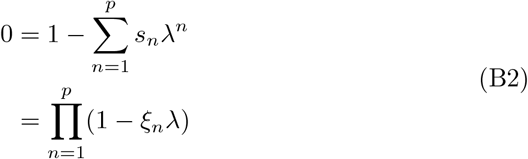

## Appendix C Prior Distributions on Parameters

### C.1 Von Mises Prior

We apply a von Mises prior to constrain the values of each oscillator frequency and prevent overfitting of large oscillations with multiple components. The von Mises prior is defined as:

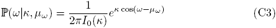

*I*_0_ is a zero-th order Bessel function. The center frequency hyperparameter *μ*_*ω*_ is initialized at the frequency of the largest pole of the AR(*p*) fit to the OSPE. The procedure for setting the concentration hyperparameter *κ* is described in Appendix G.

### C.2 Inverse Gamma Prior

In our experience, if the observation noise is underestimated, spurious oscillators may be added to the model to explain noise power at higher frequencies. To avoid this, we add an inverse gamma prior to constrain the values of the observation noise variance:

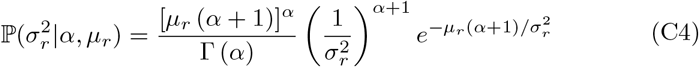

where Γ(*α*) is a gamma function. The distribution is defined on [0, ∞ ) and is concentrated around the mode, with a skewed tail going towards infinity. This spans the entire possible domain of the noise variance, which must be non-negative. *μ*_*r*_ is initialized at 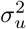. The procedure for setting the the shape hyperparameter *α* is described in Appendix G.

## Appendix D Complete Data Log Likelihood

From Shumway and Stoffer [15] we can express the complete data log likelihood as:

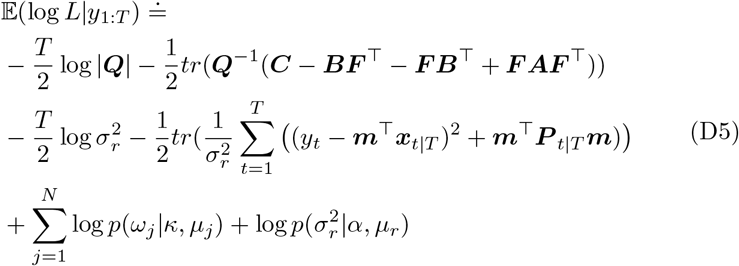

where ***Q*** ∈ ℝ^2*N*^,

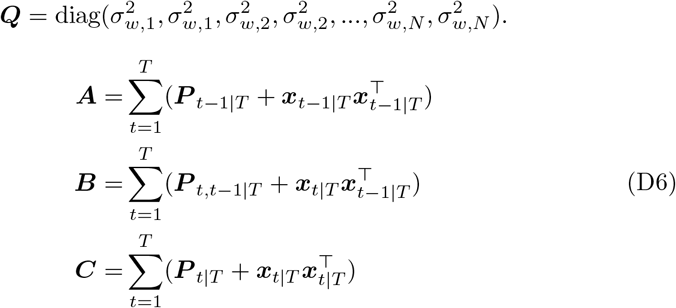

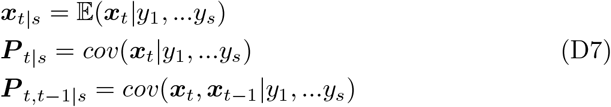

## Appendix E Update Equations for Expectation Maximization Algorithm

Since the each oscillator’s state noise is independent from the others (i.e., *Q* is a diagonal matrix) since the transition matrix *F* is block diagonal with no interaction between oscillators, each oscilaltor is independent and we can update each oscillator separately. Taking into account the priors on frequency, the update rules for each oscillator parameter and the order of updates are:

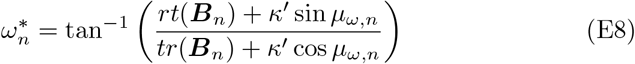

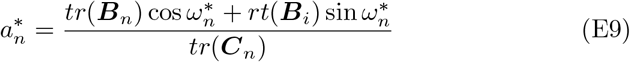

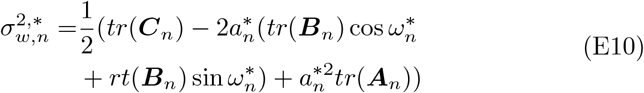

where if 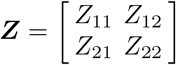, then *rt*(***Z***) = *Z*_21_ − *Z*_12_ and *tr*(***Z***) is the trace of ***Z. A***_*n*_, ***B***_*n*_, and ***C***_*n*_ correspond to the matrices in equation D6, but are specific to a single oscillation *n*. For example:

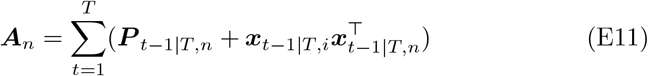

## Appendix F Using the One Step Prediction Error (OSPE) to Characterize the Unexplained Power

The one step prediction error (OSPE) depends on the difference between the one step prediction of the latent states and the smoothed latent states, which takes the current observation into account:

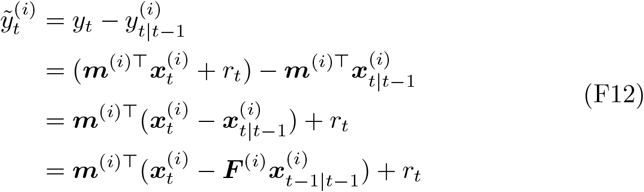

As we represent more and more of the signal (much like a greedy algorithm), the model-only prediction of the latent states. becomes closer and closer to the observation-informed prediction of the latent states.

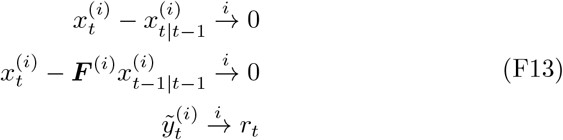

If we know that the OSPE becomes white noise, then we can make connections between this white noise and our AR(*p*) modeling of the OSPE.

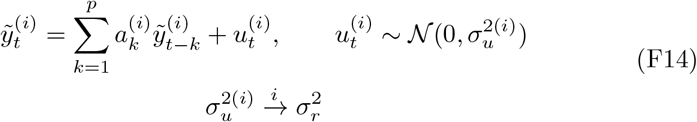

The variance of the noise driving the AR model converges to the observation noise variance.

## Appendix G Hyperparameter Selection

Here we describe the form of hyperpriors for the Von Mises and Inverse Gamma prior distributions that allow them to be insensitive to the data length.

### Von Mises Prior

The update equation for the frequency *ω* depends on the number of data points *T* and the hyperparameter *κ*:

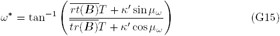

where *κ′* = *κσ*^2^/*a*. If we set

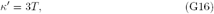

*ω*^*^ no longer depends on *T* .

### Inverse Gamma Prior

The update equation for the observation noise variance 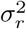 depends on the number of data points *T* and the hyperparameter *α*:

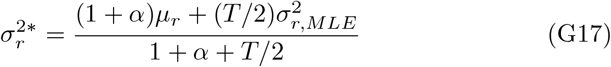

where *T* is the number of datapoints. If we set

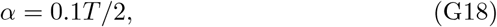

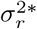 becomes insensitive to *T* for large *T* .

## Appendix H Limitations of AIC and BIC in time series data at high sampling rates

At high sampling rates, the log likelihood term increases faster than the penalty for additional model complexity. To see this, consider the OSPE or “innovations” form of the data likelihood [15]:

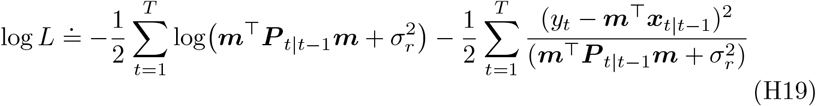

For large *T*, consider 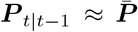, a steady-state one-step state covariance matrix. Also note that:

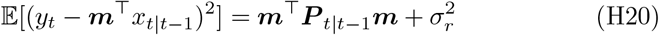

Then,

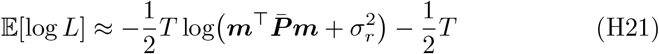

For AIC and BIC,

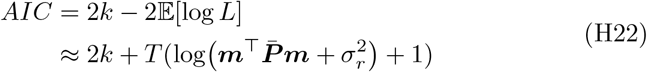

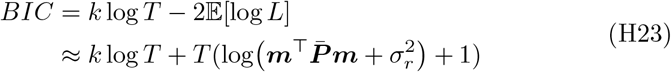

Thus when *T* is large, the log *L* term in both AIC and BIC change much faster than the model complexity terms 2*k* and *k* log *T*, respectively. (Note that log is base *e*, i.e. natural log).

When model size increases, AIC changes by:

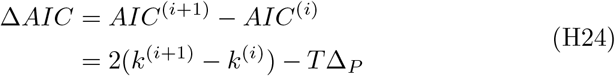

where:

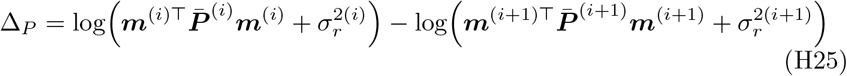

Note that if log 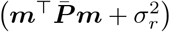 decreases, log *L* will increase.

To select adding an additional oscillator, AIC must decrease as the model order increases, meaning Δ*AIC <* 0. Using AIC, we will select adding an additional oscillation when:

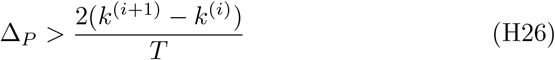

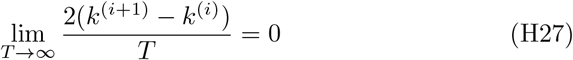

Therefore as the number of samples increases, the improvement in the log likelihood can be extremely small and still outweigh the parameter term, causing a high model order to be selected.

Likewise, when using BIC:

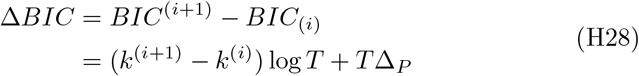

Δ*BIC <* 0 when:

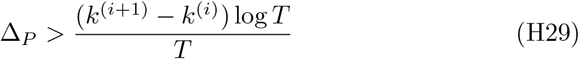

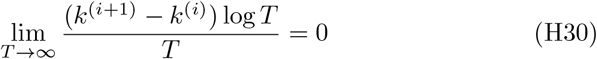

