## Supplementary Material for "An iterative search algorithm to identify oscillatory dynamics in neurophysiological time series"

#### 1 Filtering Broadband Noise

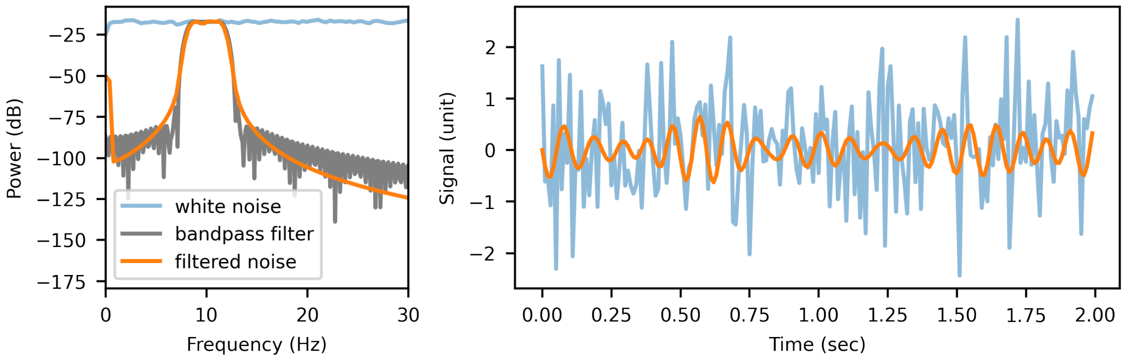


Figure 1: A bandpass filter applied to white noise produces a signal that looks like an oscillation. The power spectra were computed with Welch's method and the bandpass filter is an FIR filter of order 200 with a passband from 8 to 12 Hz. (Note that the bandpass filter spectra is actually the bandpass filter spectra multiplied by the average noise power, to better compare the shapes.)

#### 2 Data Scaling Procedure

The algorithm is intended to analyze a wide variety of signals, including EEG, MEG, ECoG, and EMG, which may be observed at different scales or in different units. To ensure that the method works well consistently, irrespective of the scale, we developed an initial normalization or scaling procedure. In particular, in simulation studies we found that the algorithm performed more consistently when the inverse gamma prior's mode and variance were comparable in scale to the observation noise. We therefore normalized the data so that each signal would have a similar observation noise variance $\approx 1$. In typical neuroscience data, oscillatory signals tend to decrease in size with increasing frequency. This implies that under our oscillator model, the observation noise will make up a larger proportion of the total observed power at higher frequencies compared to lower frequencies. We therefore obtain an initial, approximate estimate of the observation noise variance using the higher frequencies in the observed signal. After the iterative algorithm is completed and a model is chosen, we scale the model back to match the original data.

We assume that most of our signal will be in the lower frequencies and that high frequency power is mainly due to noise. We assume that a flat frequency spectrum is indicative of noise and an absence of signal and use this to determine where the frequency spectrum starts to be only due to noise. We use the average power in this flat regime to scale the observed data $y_t$ so that average power in this noise-only regime is 1. If the noise-only regime does not extend to the high frequencies, you can also choose to scale based on a smaller range of frequencies that are sufficiently flat. After the iterative algorithm is completed and a model is chosen, we scale the model back to fit the original data. The steps below describe this procedure.

1. Find power spectrum $p_y(f)$ of $y_t$ in $\mu V^2$
2. Define the flat "noise only" part of the power spectrum: either $\geq f_1$ or $\in [f_1, f_2]$. Find the average power of the "noise only" portion of the spectrum: $\overline{p}_y$.
3. Compute Scaling Factor: $c_s = \sqrt{\overline{p}_y}$.
4. Scale $y_t$ by the scaling factor: $y_{t,scaled} = y_t / c_s$
5. Apply iterative oscillator algorithm to $y_{t,scaled}$.
6. Apply inverted scaling to the fitted model to return to original scale.

$$\sigma^2_{w,j,reverted} = c_s^2 \sigma^2_{w,j,scaled}$$
$$\sigma^2_{r,reverted} = c_s^2 \sigma^2_{r,scaled}$$
$$y_{t,reverted} = c_s y_{t,scaled}$$

**Output:** Model fitting original unscaled data, $\mathcal{M}^*_{reverted}$

### 3 Von Mises Prior

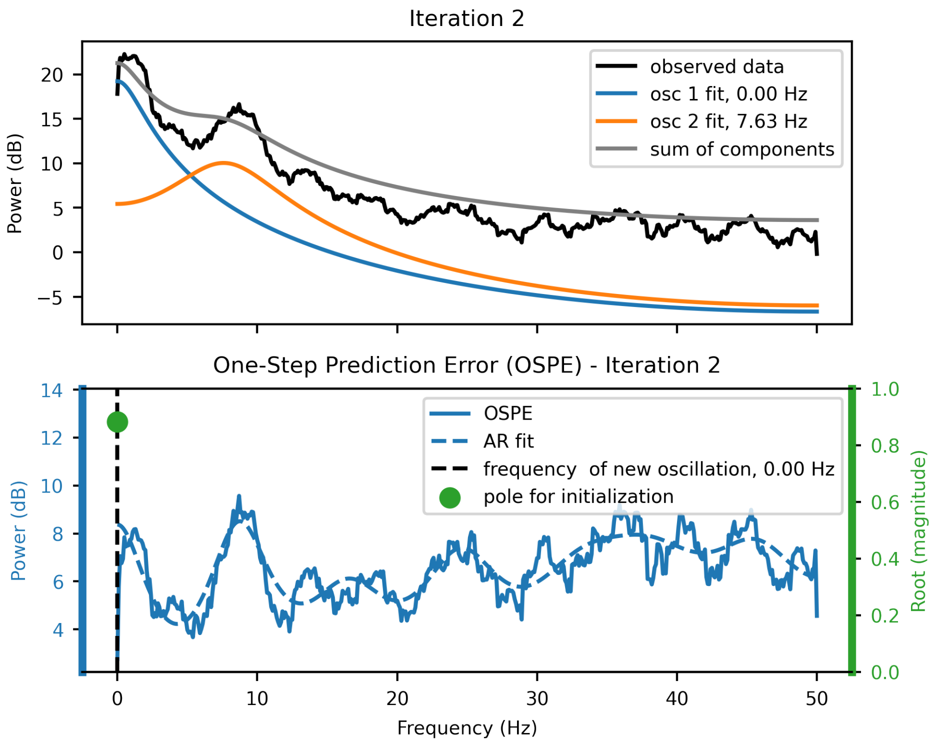


Figure 2: Top: Fitting two oscillations to the same simulated data in Main Paper Figure 1 without any priors. Bottom: One step prediction error resulting from this fitted model

#### 4 Model Parameters

For the same simulation data described in Main Paper Figure 3. The simulated data was generated based on the alpha state noise covariances listed in the legends below and the theoretical alpha spectrum based on the simulating parameters provide the theoretical alpha range (8-12 Hz) SNR, referred to in the legend as just SNR. These figures show that as the alpha oscillation becomes larger, we get more accurate measurements of the true parameters.

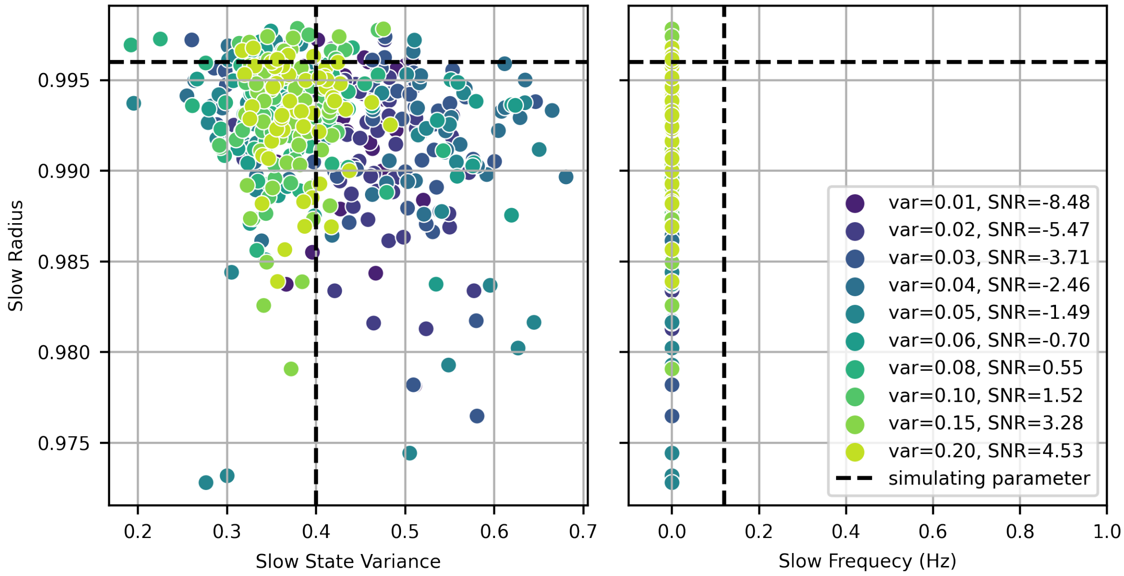


Figure 3: Parameters fitted with iOsc for simulated slow oscillations in the presence of increasing alpha (generated by increasing alpha state noise variance, "var"). All of the slow oscillations were generated with the same parameters. SNR is in dB.

The generating slow frequency of 0.12 Hz seems to be too low to distinguish from 0 Hz. Future work includes improving frequency estimation at very low frequencies. However, as shown in the next section, the model can still produce accurate oscillation estimates even in the presence of small parameter errors.

There seems to be a lower limit to the estimated alpha state variance at low SNRs, which are mostly estimated between 0.35 and 0.45. As the alpha becomes more prominent (var=0.2), the variance gets closer to the true value. The peak power of these oscillations is a function of both radius and variance, so the variance estimate benefits from a more accurate alpha radius. The alpha frequency seems to be the most robust parameter, improving at higher alpha SNRs but with almost all estimates within $\pm$ 0.5 Hz even at the lowest SNR. This due to a combination of the strong Von Mises prior on frequency and a strong model selection method. Note that only estimated models that included an alpha component are included in Figure 4.

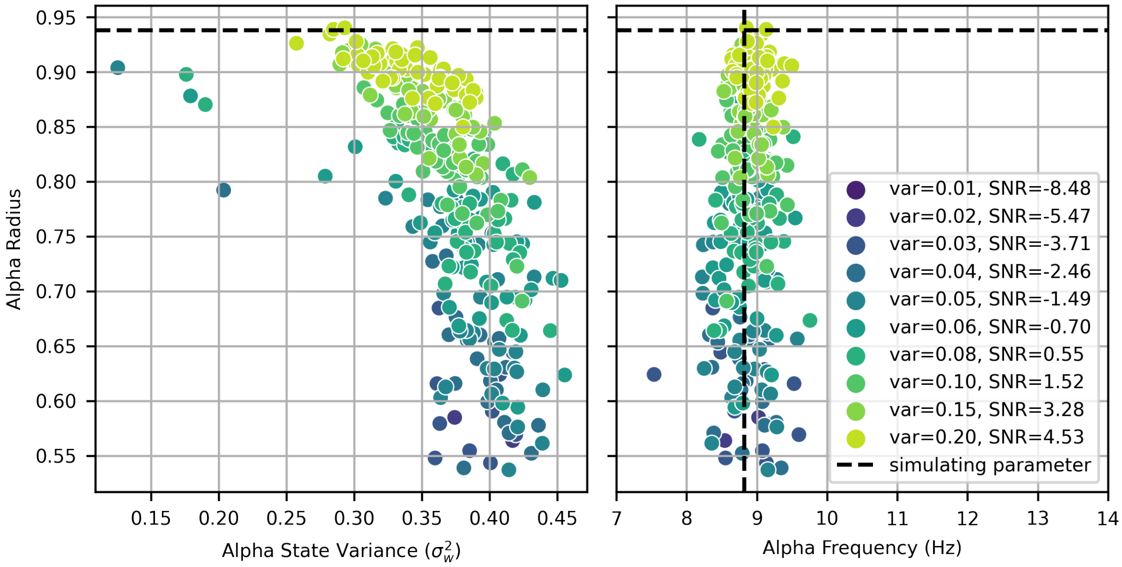


Figure 4: Parameters fitted with iOsc for simulated alpha oscillations with increasing alpha state noise variance ("var") in the presence of slow oscillations. SNR is in dB.

### 5 Spectrum Error

We see in Figure 4 that not only does our estimate of the number of oscillations get better as SNR and alpha state covariance increase, but our estimate of the simulating alpha parameters do as well. Even if there is some error in the parameter estimates, we still obtain good estimates of the time series and power spectrum because the Kalman filter and fixed-interval smoother estimate the time series based on a combination of the constraint imposed by the state space model and the observed data. Accordingly, most of the variance in the data is attributed to one of the oscillations and not the observation noise, as we saw previously in the discussion of the properties of the residuals and the OSPE. We calculate the empirical power spectrum by calculating the spectrum of the estimated alpha time series. We see in Figure 5 that the empirical power at the peak of the alpha oscillation gets more accurate as SNR increases, and the model tends to underestimate peak frequency at low SNR.

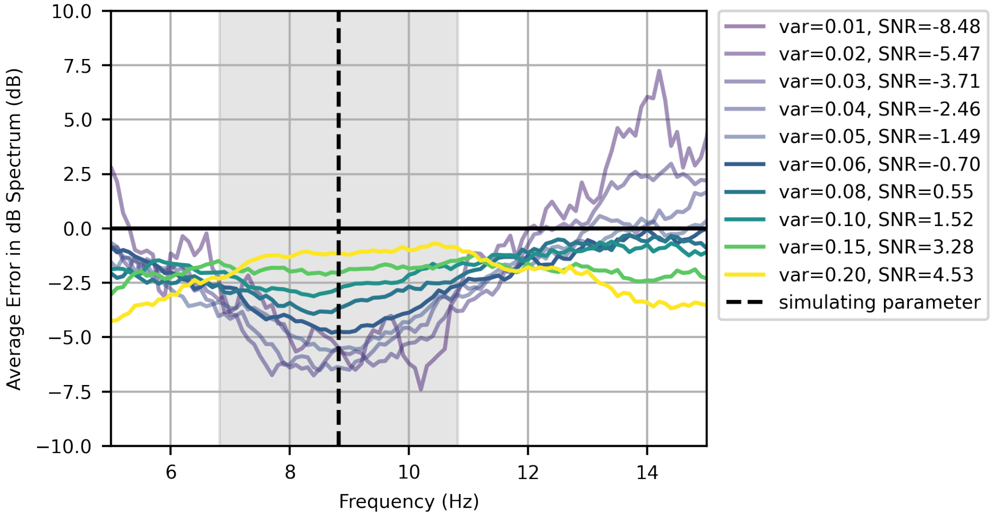


Figure 5: The average difference between the true empirical spectrum of the simulated alpha (in dB) and the estimated empirical spectrum (in dB)

#### 6 Relationship between Alpha SNR and Noise

Although the alpha state covariance simulating parameter allows us to manipulate the SNR of alpha, the SNR can vary based on the specific data simulated. We see in Figure 6 that the SNR may overlap across different simulated noise covariances, but it is tightly correlated with alpha noise covariance and the number of oscillations selected. The SNR in Figure 6 is the ratio of alpha to non-alpha power in the alpha band (8-12 Hz), showing that we start to identify a second oscillator more often than chance when the alpha SNR is slightly below -2.5 dB.

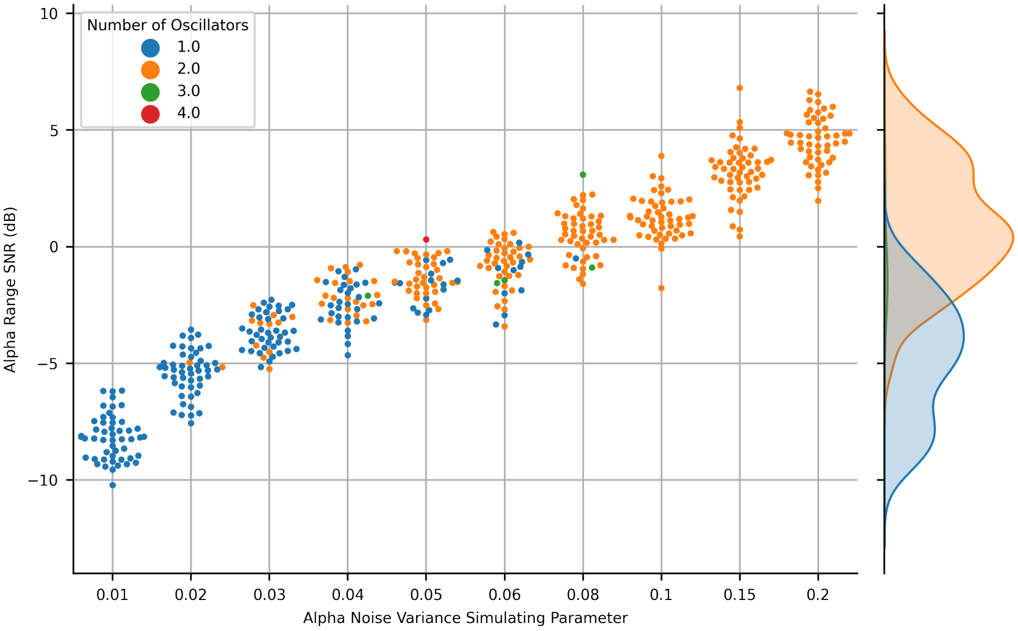


Figure 6: Left: Empirical alpha range SNR of each simulated data set vs simulating variance parameter. Right: Distribution of alpha range SNR dependent on how many oscillators were selected.

Figure 7 shows that there is a very clear distinction between selecting one and two oscillations that depends on the relationship between the not only on the alpha range SNR but also the estimated observation noise variance. When the observation noise is estimated by the model at a higher level, it does not select an alpha until higher alpha range SNRs. This is consistent with our observation that accurate estimation of the observation noise variance is crucial for accurate model selection (i.e. selecting the correct number of oscillators). When an oscillation is small, if the observation noise estimate is large the oscillation can be attributed to noise.

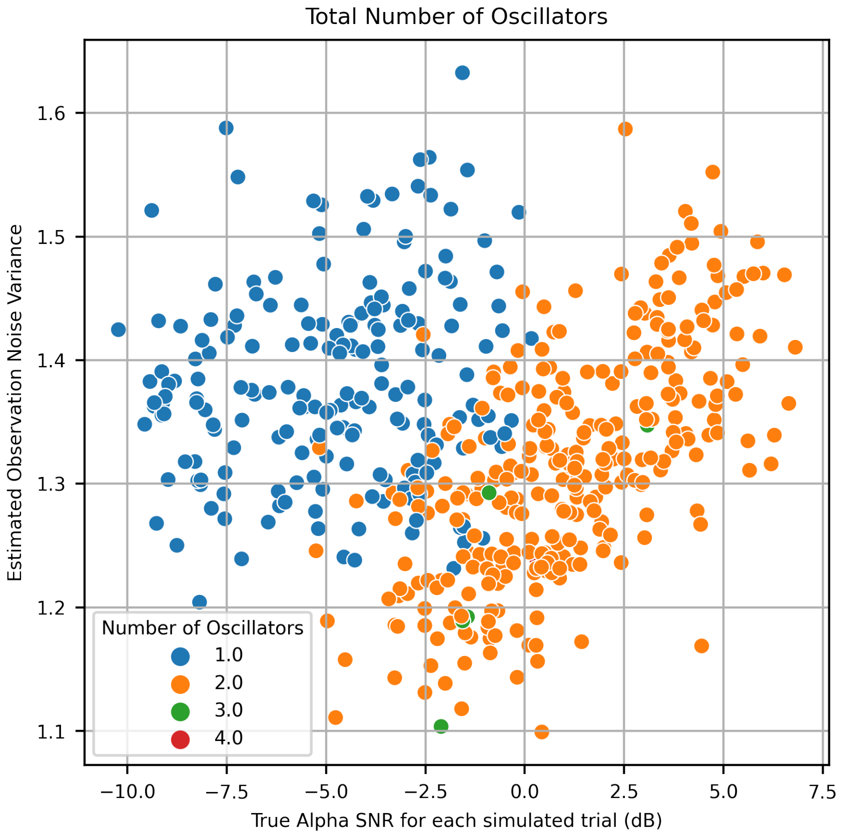


Figure 7: Relationship between empirical alpha range SNR (from the true simulated alpha) and the observation noise variance estimated by iOsc

### 7 Comparing Methods: A Simulation Case Study

One example simulated data set from the highest SNR level described previously (theoretical alpha range SNR= 4.53 dB) showing common pitfalls of thee FOOOF and Matsuda and Komaki methods.

#### 7.1 iOsc Model

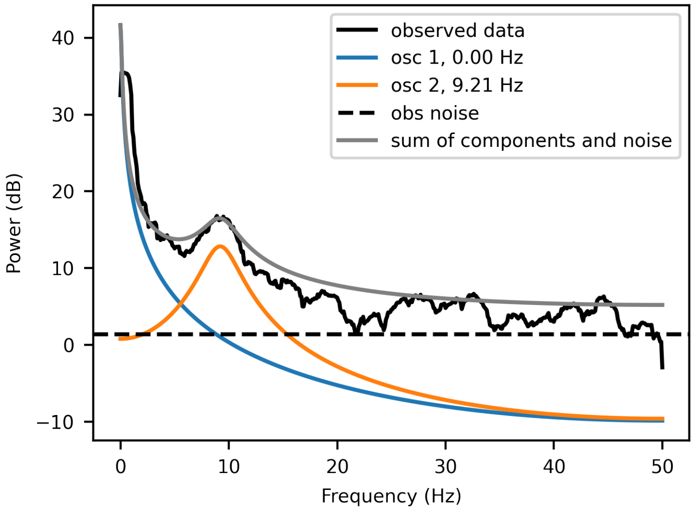


Figure 8: Subject from highest alpha SNR from Main Paper Results: Simulation

#### 7.2 FOOOF

The FOOOF method tends to add extra oscillations to explain any deviation from the aperiodic fit.

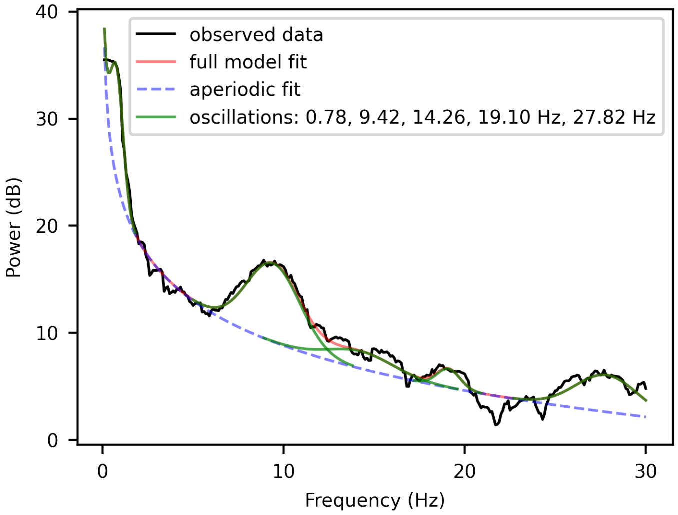


Figure 9: FOOOF fit gaussians to the multitaper spectrum of the simulated data (SR=2 Hz). Center frequencies listed in the legend.

##### 7.3 Matsuda and Komaki

The Matsuda and Komaki method tends to add more oscillations than necessary, both near the frequency of a true oscillation (alpha) and in higher frequencies, possibly due to ill-fitting observation noise variance estimates. Some of these oscillations are decreased to very low power, so we consider them to be negligible, but there are still extra oscillators that could affect the fits of the true oscillators.

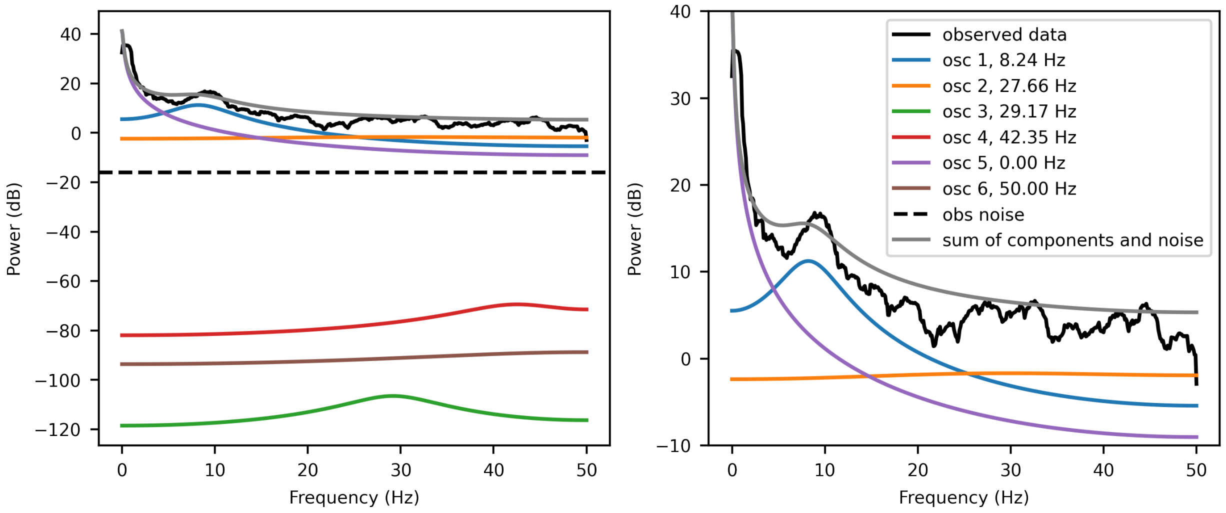


Figure 10: Left: Matsuda and Komaki model fitted to simulated data. Right: Same fitted model, zoomed in on the observed data section.

###### 7.3.1 Implementation details

The Matsuda and Komaki method initializes parameters based on the outcome of an AR model fit to the observed data. They consider a range of AR orders based on a user-defined number of expected oscillations. To avoid this definition by the user, we fit AR models up to order 30 and then use AIC, as they describe, to determine the optimal number of oscillators.

Once the oscillator frequencies and radii are determined by the selected AR model, Matsuda and Komaki describe two methods to calculate the noise variance for each oscillation. The simplest method relies on solving a matrix equation, matching the sum of the oscillation power spectra to the sample spectrum. This produces several problems. The sample spectrum has very high variance, which can produce anomalously low variances, possibly causing some of the poor fit. If the AR fit produces several positive real roots, then not all center frequencies are unique and the matrix, which is the power spectrum of each oscillation evaluated at the center frequency of each oscillation, becomes singular. Even when it does not become singular, it can produce negative variances, is incompatible with variance. To account for both the possible singularity of the matrix we need to invert and the possible negative variances, we solve the matrix equation using a least squares fit with a soft L1 penalty, and restrict the variance to nonnegative values. The initialized variance is then computed as described in Appendix 2 of Matsuda and Komaki, where it is the evaluated values of the matrix equation scaled by the fitted observation noise variance.

Using these initial parameters, we use an expectation maximization algorithm without priors to fit the model parameters. Due to the lack of priors, the oscillations can shift toward the high power oscillations, as described in our main paper. Since the initial observation noise is extremely low, the state noise variances become very high and this also accounts for some of the errors in fitting. Also note that although the AR models described by Matsuda and Komaki identify single real roots, which describe an AR(1) process, their initialization process and oscillator structure inserts a duplicate real root, creating an ARMA(2,1) process and changing the power spectrum from the originally defined one.

### 8 Comparing Methods: An EEG Case Study

In this case we do not know the ground truth, but we do expect to see a slow wave and an alpha oscillation in occipital electrodes with eyes closed at rest [26-28] as described in the main paper. The following section shows how the errors seen in the previous section impact our results on real data.

#### 8.1 iOsc Model

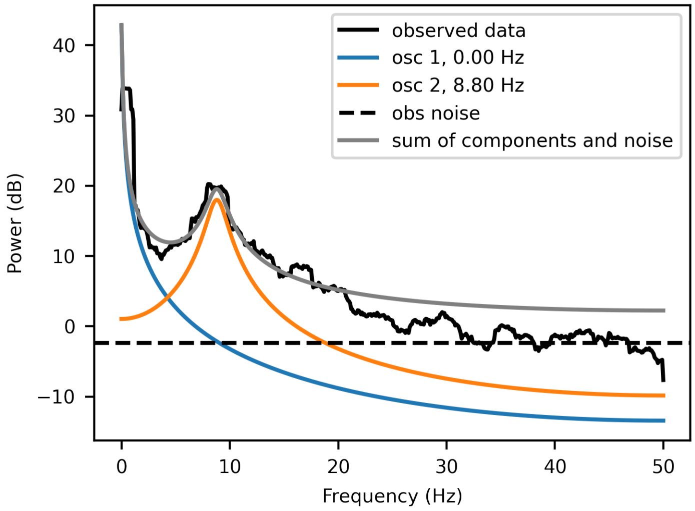


Figure 11: Subject 1 from Main Paper Results: Resting EEG section 4. Reproduced for comparison with sections 8.2 and 8.3

#### 8.2 FOOOF

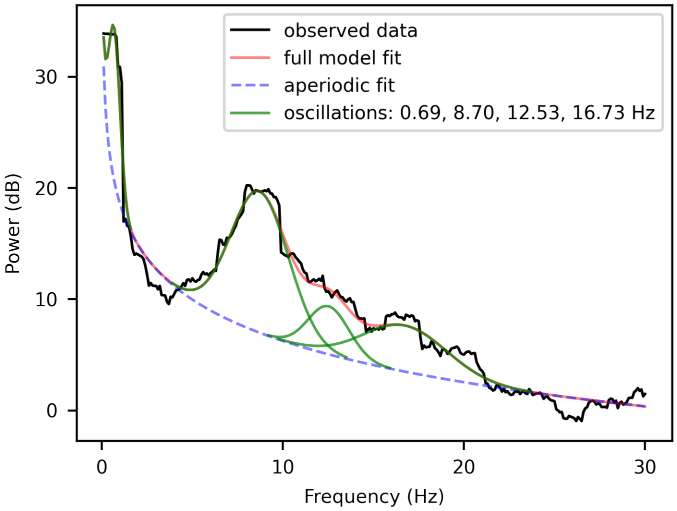


Figure 12: FOOOF applied to resting state eeg data Subject 1

##### 8.3 Matsuda and Komaki

The Matsuda and Komaki method has fitted the observed data spectrum reasonably well, however it includes many superfluous oscillators and notably includes two alpha oscillators to represent what is clearly a single oscillation, which can lead to poor fitted parameters for each.

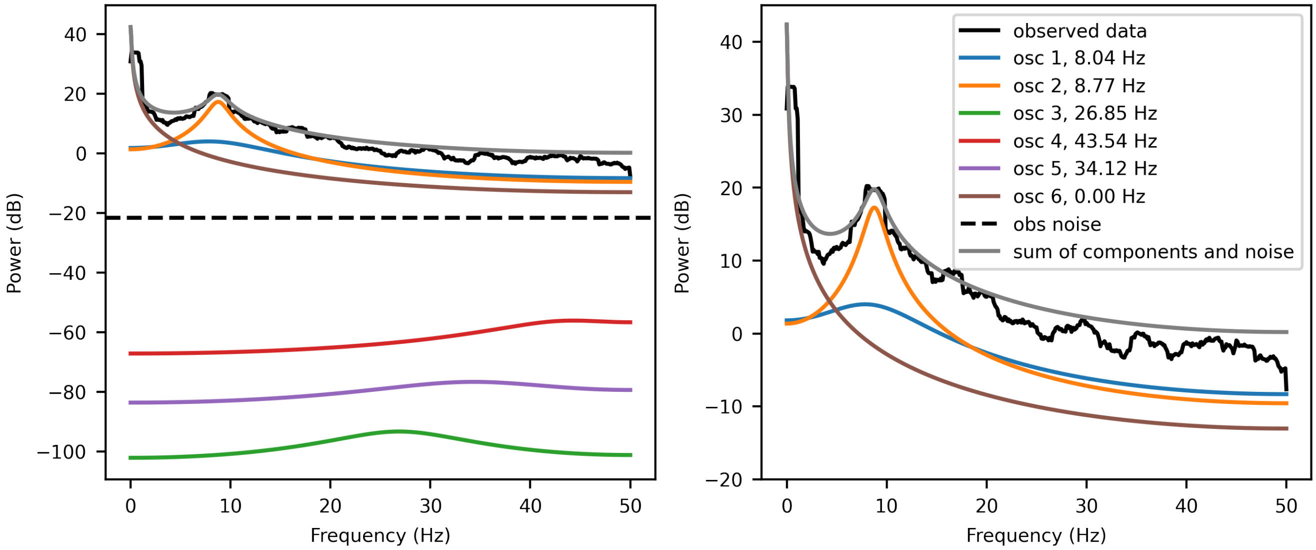


Figure 13: Left: Matsuda and Komaki model fitted to resting state eeg data Subject 1. Right: Same fitted model, zoomed in on the observed data section.

### 9 MoCA Model Fitting

When fitting a nested model the MoCA data, we see that the logistic model provides a sharper cutoff in the case of iOsc and the iOsc data is closer to the linear fit than in the case of FOOOF. This generalized linear model was fit using R on the frequency values and an indicator of the presence of alpha. For FOOOF, the coefficent on frequency was 0.7297 and 5.4906 on the indicator with an intercept of 19.7094. For iOsc, the cofficient on frequency was 0.9812 and 6.7293 on the indicator, with an intercept of 17.985.

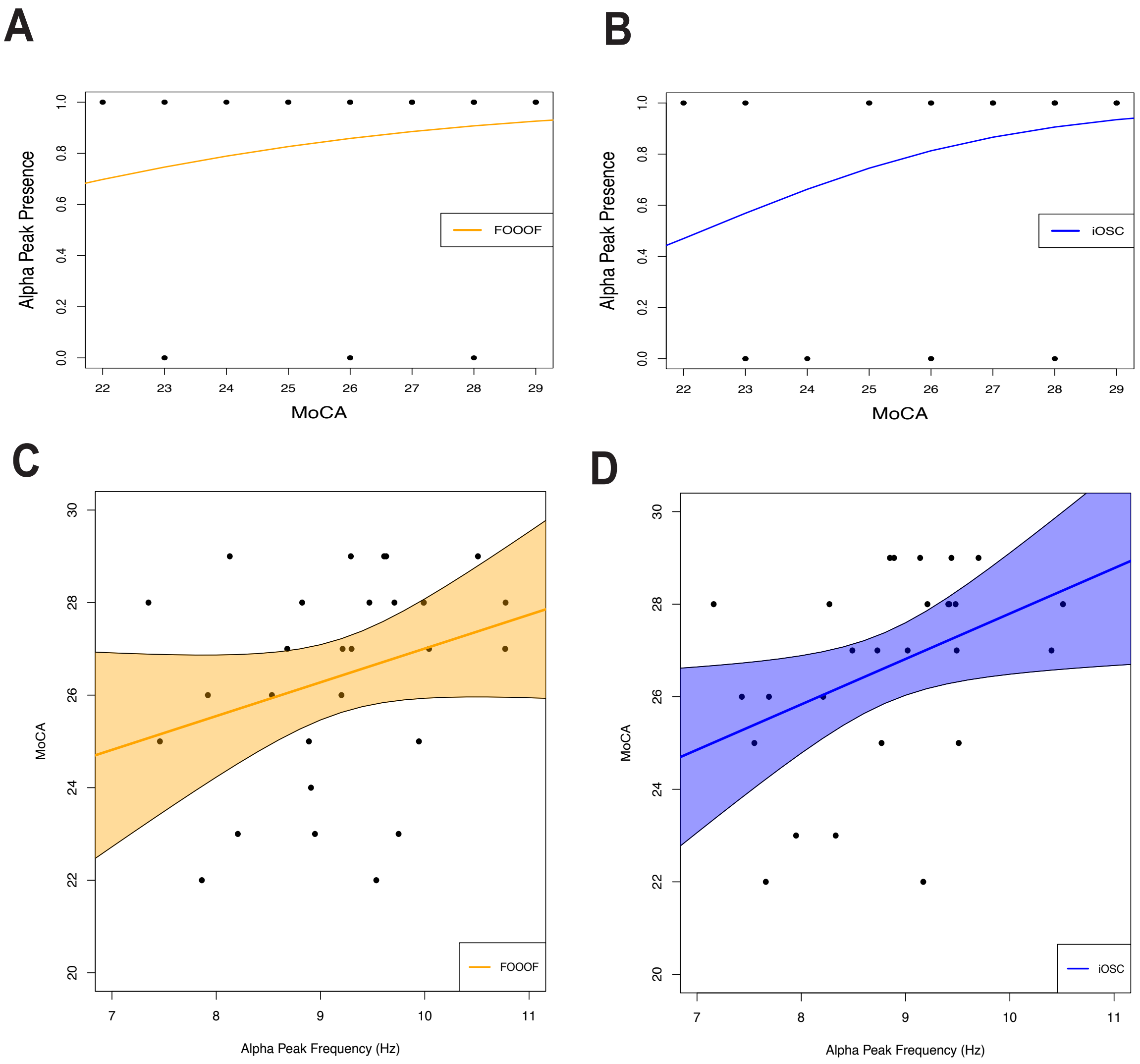


Figure 14: Nested model showing the relationship beween MoCA and alpha peak frequency. Top row: logistic model to include presence of alpha. Bottom row: linear model to evaluate correlation between alpha peak frequency and MoCA

### 10 A Note on iOsc Model Fitting

One major goal of this method was to produce an algorithm that required minimal user intervention, if any. We recommend starting with the algorithm as is, but in the case of poor fitting, we suggest the following alterations:

1. If the pole initialized from the one-step prediction (represented by the green circle in Figure 2, bottom panel) is between two oscillations, causing poor fitting of this oscillation as it attempts to explain multiple oscillations, we recommend increasing the order of the AR model used to approximate the OSPE. Increase in increments of two, which will allow additional pairs of complex poles.

2. Conversely to point 1, if the order of the AR model is too high then multiple pairs of roots will be attributed to the same oscillation, diluting the strength needed for each of them and possibly leading to none of them being selected as the strongest root in the iterative process to initialize the next oscillation, even though together they describe the strongest oscillation. This can be identified using the innovations plot with all of the AR roots plotted. In this case we recommend decreasing the AR order in increments of 2, to decrease the number of pairs of complex poles.

3. If the initialization of the additional oscillations describes a single oscillation well, but the fitting of this oscillation attempts to explain multiple oscillations and causes poor fitting, we recommend increasing the concentration hyperparameter in the Von Mises prior. This will increase the weight on the initial frequency and stop the oscillation from shifting to explain other oscillations.

4. If the model does not choose the correct number of oscillations, we recommend looking at all fitted models and selecting the best fitting model based on other selection criteria or using your best judgement. You can also choose a subset of well-fitted oscillations and run the kalman filter to estimate oscillations using those fitted parameters.

5. Note that this algorithm assumes a stationary signal, and therefore stationary parameters. Although the Kalman filtering allows some flexibility in this requirement, enabling the model to work on some time-varying signal, the success of the method depends on the strength and duration of the signal components. The weaker and more brief the time-varying component is, the more poorly the model will capture it, if it does at all. We recommend decreasing the length of your window until you have a more stationary signal.

This algorithm is designed to fit well automatically in most situations, but there will still be some data sets where it does not fit well without intervention.
